# Physiological stress drives the emergence of a *Salmonella* subpopulation through ribosomal RNA regulation

**DOI:** 10.1101/2023.03.02.530801

**Authors:** Camilla Ciolli Mattioli, Kfir Eisner, Aviel Rosenbaum, Mengyu Wang, Ariel Amir, Ido Golding, Roi Avraham

**Affiliations:** Department of Immunology and Regenerative Biology, Weizmann Institute of Science, Rehovot, Israel; Department of Physics of Complex Systems, Weizmann Institute of Science, Rehovot, Israel; Department of Physics, University of Illinois at Urbana-Champaign, Urbana, IL, USA

## Abstract

Bacteria undergo cycles of growth and starvation, to which they must adapt swiftly. One important strategy for adjusting growth rates relies on ribosomal levels. While high ribosomal levels are required for fast growth, their dynamics during starvation remain unclear. Here, we analyzed ribosomal RNA (rRNA) content of individual *Salmonella* cells using Fluorescence In-Situ Hybridization (rRNA-FISH). During the transition from exponential to stationary phase we measured a dramatic decrease in rRNA numbers only in a subpopulation, resulting in a bimodal distribution of cells with high and low rRNA content. We showed that the two subpopulations are phenotypically distinct when subjected to nutritional upshifts. Using a transposon screen coupled with rRNA-FISH, we identified two mutants acting on rRNA transcription shutdown and degradation, that abolished the formation of the subpopulation with low rRNA content. Our work suggests that *Salmonella* employs a bet-hedging strategy in regulating ribosomal levels that may be beneficial for survival.

## Introduction

Bacterial cells are faced with the constant challenge of optimizing resource allocation in response to changing environments. Inside the host gut, enteric bacteria such as *Salmonella enterica* serovar Typhimurium (S.Tm) undergo alternate growth and starvation periods dictated by the feeding cycles, commonly referred to as “feast and famine”. Furthermore, during intracellular infection of macrophages, *Salmonella enterica* resides in a modified endosomal compartment restricted in nutrients [1]. During starvation, an important survival strategy consists in the breakdown of ribosomes [2, 3], constituting a large fraction of cellular mass. This process allows the release and recycling of essential micromolecules, replenishing the pool of amino acids and ribonucleotides.

In rapidly dividing bacteria, protein synthesis is maximized by employing all ribosomes in full capacity [4]. Under these conditions, ribosomal RNA (rRNA) is protected from degradation by the engaged monosomes [5]. In contrast to the high stability exhibited during exponential growth [3, 6], rRNA has been reported to degrade in response to several types of starvation [7–12]. The decrease in translation that occurs during starvation results in the disassembly of monosomes into free ribosome subunits, allowing rRNA endonucleolytic cleavage and ultimately, ribosome breakdown [5]. A few reports found RNase II and RNase R exoribonucleases to be involved in the hydrolysis of the rRNA fragments [13] generated by an endoribonuclease, yet to be identified conclusively [14, 15]. Alongside ribosomal degradation, the alarmone molecule (p)ppGpp is synthesized following starvation [16–21]. (p)ppGpp accumulation has been associated with the repression of transcription of rRNA and other genes [22–24], along with the induction of hibernation factors [25–29], known to form a complex with nontranslating monosomes to protect them from degradation [15, 30]. While ribosomal degradation is considered an adaptation strategy during starvation, we still have a limited understanding of its dynamics, underlying regulation, and potential impact on bacterial fitness.

Here, we studied the rRNA content of individual *S*.Tm cells using Fluorescence In Situ Hybridization (rRNA-FISH) during the transition from exponential to stationary phase. At the population level, we observed an asynchronous decrease in rRNA that resulted in the formation of two subpopulations, characterized by low and high rRNA content. We further analyzed the phenotypes associated with the two subpopulations, and showed a fitness advantage during nutritional upshifts for the subpopulation of bacteria maintaining high ribosomal levels. Finally, using a transposon screen coupled with rRNA-FISH and flow cytometry [31], we identified DksA and RNase I as molecular drivers in the formation of the subpopulation with low ribosomal content acting via rRNA transcriptional shutdown and active rRNA degradation. Our results provide compelling insights into the strategies that bacterial pathogens employ to adapt and survive to changing environments.

## Results

### *S*.Tm displayed heterogeneous levels of ribosomes in the transition from exponential growth to stationary phase

To quantify rRNA levels in single cells, we optimized an rRNA-FISH approach for bacterial cells [32]. rRNA-FISH has the advantage of directly assessing RNA levels, while fluorescent reporters based on fusions with ribosomal proteins may alter functionality [33] and half-life [34]. We used spectrally distinct fluorescent probes against 16S and 23S rRNA for measuring *S*.Tm rRNA corresponding to ribosomes (Extended Data Fig. 1a). The probes were designed on a highly conserved region of the rRNA operon to anneal on any of the seven copies present in the *S*.Tm genome (Extended Data Fig. 1b). To confirm that the 16S signal corresponded to rRNA loaded into ribosomes, we used two different probes and imaged mature 16S and premature 16S rRNA separately. In bacteria with a low nucleoid-to-cytoplasm ratio such as *S*.Tm, ribosomes are enriched in the nucleoid-free region of the cytoplasm [35], while the 5’ leader sequence of premature newly transcribed rRNA is retained in the nucleoid, where it is rapidly processed by cleavage after transcription [36]. Accordingly, we observed a spatial segregation of mature 16S rRNA from the nucleoid and a spatial overlap of the premature 16S 5’ leader sequence (pre-16S) with the DAPI-stained nucleoid (Extended Data Fig. 1c,d), suggesting that the 16S probe localization corresponds to rRNA loaded into ribosomes. Using microscopy (Extended Data Fig. 1e), we analyzed the mean fluorescence intensity per pixel within the area of individual cells and, using flow cytometry (Extended Data Fig. 1f), the total fluorescence intensity per cell of 16S and 23S rRNA, belonging to the 30S or 50S ribosomal subunits, respectively. We observed a high correlation (R = 0.73 – 0.98) in fluorescence intensity between the two probes, further indicating that they are associated with assembled ribosomes (Extended Data Fig. 1e,f). From the imaging data, we measured several parameters encompassing cell length, cell area, and number of chromosomes (Extended Data Fig. 1g). As expected, cells that underwent exponential cell division were larger and longer (Extended Data Fig. 1g, upper and middle panel), and accordingly, in these cells the number of chromosomes was greater than one (Extended Data Fig. 1g, lower panel). Furthermore, we excluded the presence of dead or damaged cells that could potentially cause probe binding artifacts by assessing cellular permeability with propidium iodide (Extended Data Fig. 1h). Finally, we compared averaged 16S rRNA-FISH measurements to bulk RNA / protein ratios (*r*) obtained from growth in different media [4], indicative of ribosome levels. We observed a high correlation (R = 0.99) between 16S rRNA-FISH and RNA / protein ratios, confirming the performance of 16S rRNA-FISH (Extended Data Fig. 1i).

To follow rRNA dynamics in individual cells at different growth stages, a preculture of *S*.Tm was inoculated into fresh Luria Bertani (LB) medium. At the indicated time points, representing different stages of growth (indicated by the colored areas in Fig. 1a), *S*.Tm was subjected to rRNA-FISH and imaging. By tracking the mean fluorescence intensity per pixel within the area of individual cells, we observed that 16S rRNA-FISH signal increased rapidly soon after cells exited lag phase and reached a high level that was uniform across the population during exponential growth. Interestingly, during the transition to stationary phase, two subpopulations of cells became apparent, characterized by high or low ribosomal levels (Fig. 1b,c). The fraction of 16S^*high*^ cells decreased as bacteria entered stationary phase, together with an increase in the fraction of 16S^*low*^ cells. A small proportion of cells maintained 16S^*high*^ levels across several time points during the transition to stationary phase (1-6%).

**Fig. 1:**
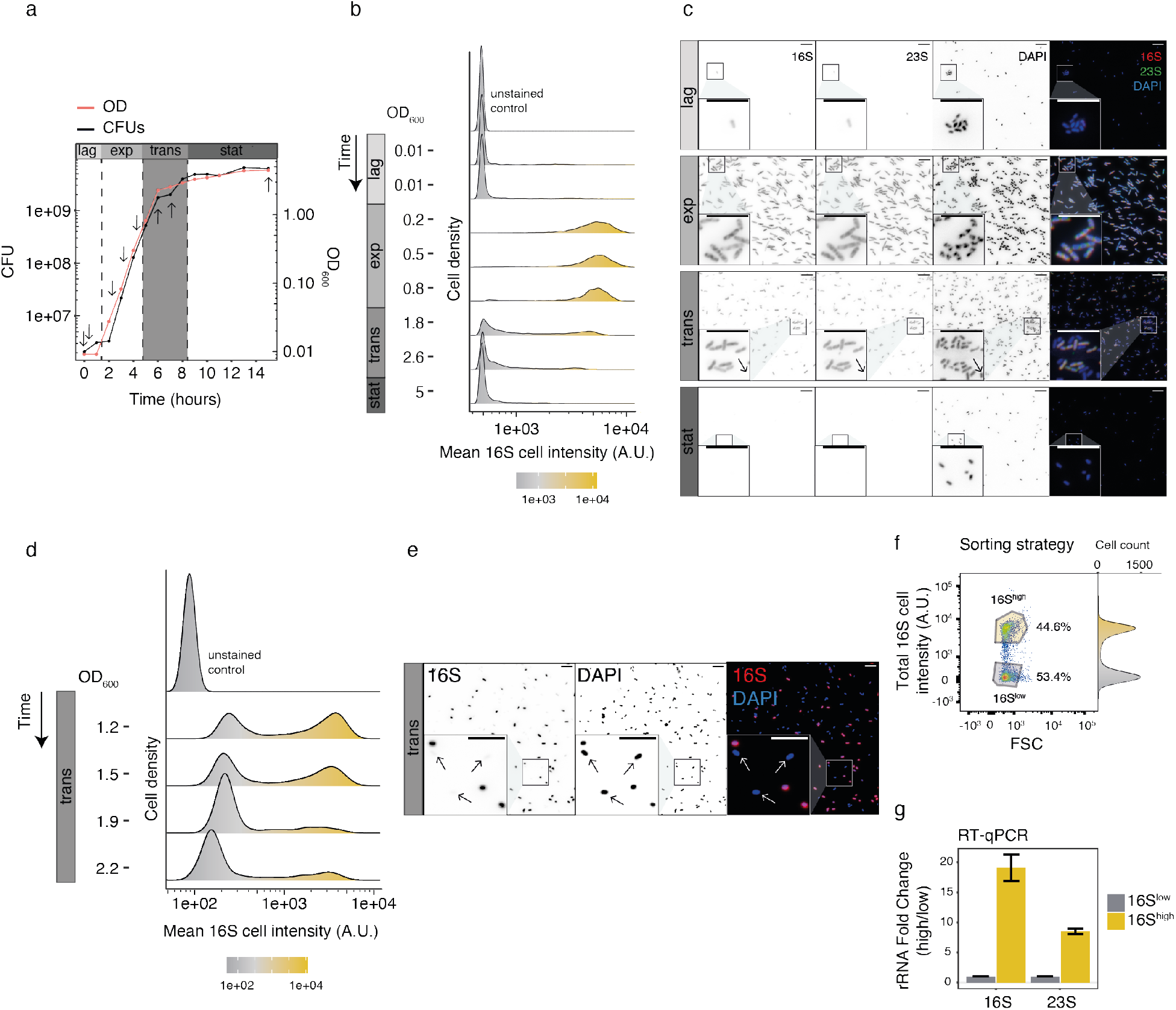
S.Tm displayed heterogeneous levels of ribosomes in the transition from exponential growth to stationary phase. **a**, Growth curve of *S*.Tm was measured by CFUs/ml (in black) and OD_600_ (in light red) during 16 hours in LB medium. Estimated growth phases are indicated on the top (lag; exp, exponential; trans, transition; stat, stationary). The grey area corresponds to the transition from exponential to stationary phase. Arrows indicate the time points assayed in Fig. 1b. **b**, Ribosomal levels decrease upon transition from exponential growth to stationary phase. Density histograms show 16S rRNA-FISH mean fluorescence intensity (total fluorescence divided by the number of pixels per cell) of individual *S*.Tm cells measured from microscopy images (n = 967 – 6,025 cells). Each sample OD_600_ is indicated on the left. Non-stained bacteria are shown (unstained control). **c**, Representative large field and close-up microscopy images of four growth phases (lag, exponential, transition, and stationary) from Fig. 1b are shown. 16S rRNA-FISH, red; 23S rRNA-FISH, green; DAPI, blue. Single channels are shown in black. Arrows indicate cells with low levels of ribosomes (16S^*low*^ cells). Scale bar, 10 *μ*m. **d**, Detailed measurements in the transition phase indicate sustained bimodality in 16S rRNA signal. Density histograms show 16S rRNA-FISH mean fluorescence intensity of individual *S*.Tm cells measured from microscopy images during to the transition from exponential growth to stationary phase (n = 554 – 2,027 cells). Non-stained bacteria (unstained control) are shown. Each sample OD_600_ is indicated on the left. **e**, Representative large field and close-up microscopy images of the transition phase from Fig. 1d are shown. 16S rRNA-FISH, red; DAPI, blue. Single channels are shown in black. Arrows indicate cells with low levels of ribosomes (16S^*low*^ cells). Scale bar, 10 *μ*m. **f**, Flow cytometry analysis indicating the gates used for sorting 16S^*low*^ and 16S^*high*^ subpopulations based on 16S rRNA-FISH total fluorescence intensity (per cell) and forward scatter (FSC) areas. Histogram of the data in the y axis is provided. **g**, RT-qPCR validates the difference in ribosomal content between 16S^*low*^ (in grey) and 16S^*high*^ (in yellow) cells. Values are shown as fold change using 16S^*low*^ as control. Error bars indicate means ± SEM of three technical replicates.

Reduction in rRNA abundance in the transition phase has previously been described at the bulk level in *E. coli* [3] and *Salmonella* [37]. However, due to the limitation of bulk measurements, the bimodality we report here in single cells was not observed and its implications were not investigated. To follow the appearance of the bimodality in the transition phase, we performed a finer time-course experiment, sampling every 20 minutes during the transition phase (Fig. 1d). We measured a bimodal distribution of 16S rRNA abundance that persisted longer than an hour. The 16S^*high*^ subpopulation showed a 14-fold increase in 16S rRNA-FISH signal compared to the 16S^*low*^ subpopulation. We validated our rRNA-FISH measurements using RT-qPCR to exclude the possibility that the reduction in rRNA-FISH fluorescence was due to cellular permeability, rRNA modifications, or ribosomal conformational changes affecting probe accessibility and binding. We analyzed 16S rRNA-FISH fluorescence levels of *S*.Tm in the transition phase by flow cytometry (Fig. 1f), and sorted 16S^*low*^ and 16S^*high*^ subpopulations using fluorescence activated cell sorting (FACS), followed by RNA extraction and RT-qPCR using primers targeting the 16S or 23S genes. Potential gDNA contamination was excluded by using a no reverse transcriptase control (NRT), where comparable background levels were measured in the two subpopulations (Extended Data Fig. 1j). Corroborating our rRNA-FISH results, the 16S^*high*^ sample showed in average 19- and 8.5-fold increase over the 16S^*low*^ subpopulation for 16S and 23S rRNA, respectively (Fig. 1g), in line with previous reports demonstrating a 90% reduction of rRNA in *Salmonella* upon entry into stationary phase [37].

### Ribosomal bimodality was conserved among *Salmonella* strains and also observed during intracellular infection

Next, we sought to test whether the ribosomal bimodality displayed at the transition phase is unique to specific *Salmonella* strains or is a general phenomenon. Three *S*.Tm strains (14028S, D23580, and S4/74) showed similar behavior, where both 16S^*high*^ and 16S^*low*^ sub-populations were present (Fig. 2a and Extended Data Fig. 2a,b). Unlike *S*.Tm, we did not observe a clear bimodal state in *E. coli* nor *S. aureus* (Fig. 2a). We performed a finer time course experiment in *E. coli*, and measured its 16S rRNA levels. We observed a heterogeneous 16S rRNA decrease during the transition phase that stabilized to low, uniform levels in stationary phase (Fig. 2b). These findings confirm that both *S*.Tm and *E. coli* regulate their ribosomal content by reducing it in response to nutrient scarcity, as it has been previously reported [3]. Unlike previous reports, using single cell measurements, we observed a rapid formation of bimodality in *S*.Tm ribosomal levels that separated the population into two distinct subgroups characterized by 16S^*high*^ and 16S^*low*^ levels, revealing an underlying heterogeneity in the process of ribosomal decrease.

**Fig. 2:**
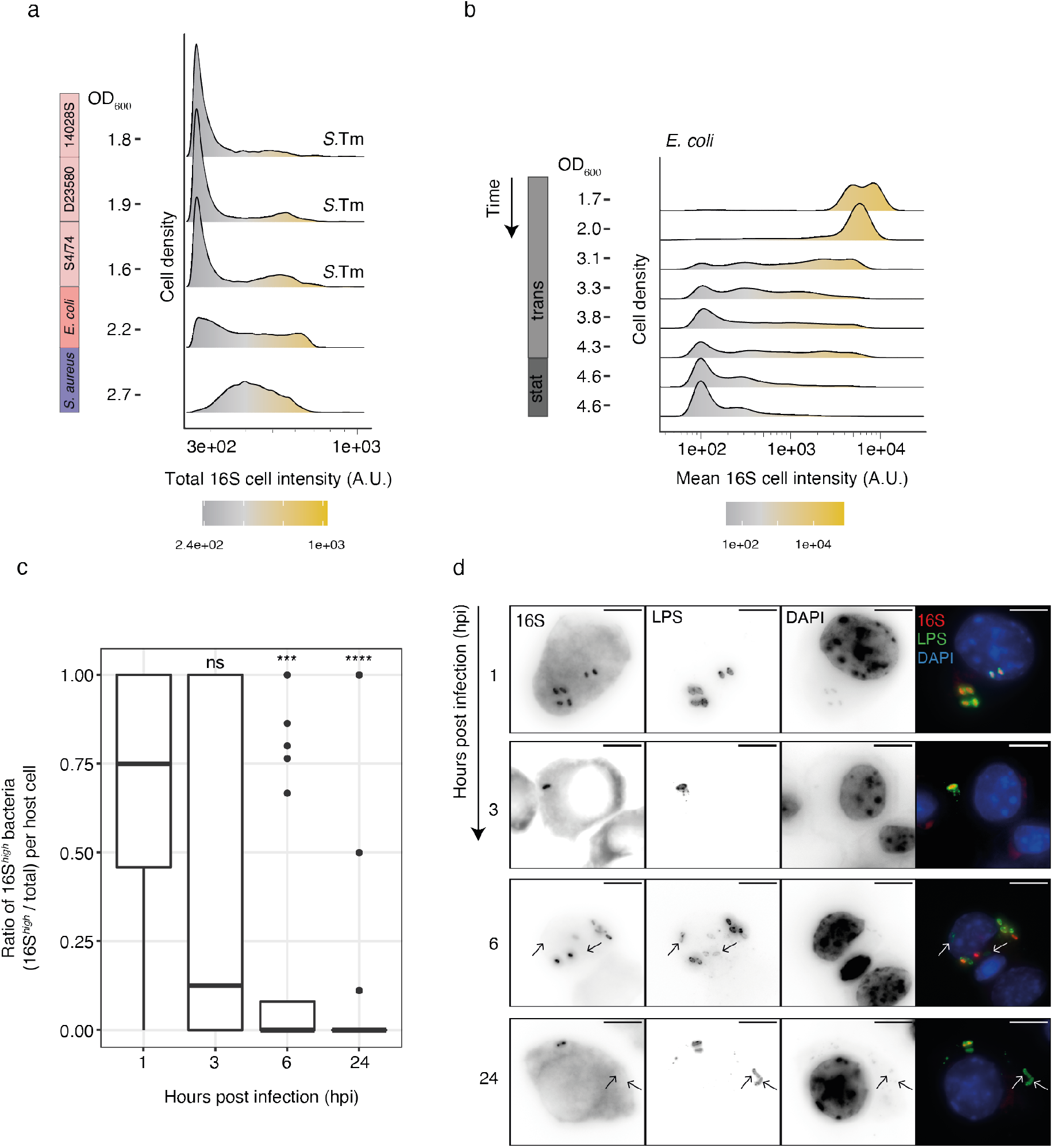
Ribosomal bimodality was conserved among *Salmonella* strains and also observed during intracellular infection. **a**, Ribosomal bimodality is conserved across several *S*.Tm strains. Density histograms show 16S rRNA-FISH total fluorescent intensity (per cell) of individual *S*.Tm cells measured by flow cytometry, from the indicated bacterial strains (n = 11,208 – 31,407 cells). Each sample OD_600_ is indicated on the left. **b**, Density histograms of 16S rRNA-FISH mean fluorescence intensity of individual *E. coli* K12-MG1655 cells measured from microscopy images (n = 977 – 3,114 cells). Each sample OD_600_ is indicated on the left. **c**, During infection in macrophages, 16S rRNA levels decrease in *S*.Tm. The fraction of 16S^*high*^ bacteria over the total measured per infected J77A-1 macrophage cell is shown. Samples were taken at 1, 3, 6, and 24 hours post infection (n = 35 – 89 host cells, n = 112 – 526 bacteria). Unpaired t-test: ns, *P* > 0.05; ***, *P* <0.001; ****, *P* <0.0001 (reference group: 1 hpi). **d**, Representative microscopy images of the samples from Fig. 2c are shown. 16S rRNA-FISH, red; LPS, green; DAPI, blue. Single channels are shown in black. Arrows indicate 16S^*low*^ bacteria. Scale bar, 10 *μ*m.

To further investigate how the observed bimodal ribosomal distribution can arise, we constructed a theoretical framework consisting of two models (see Supplementary Notes 1 for detailed explanation). The first model consists in a dilution-based process following rRNA transcription shut down, where ribosome concentration is halved with each population doubling. The second model expands on the first by including an rRNA degradation mechanism. Both models assume that at specific point in time, bacterial cells detect a depletion in nutrients in the environment, and gradually switch from a state of exponential growth to a state where they shut down rRNA transcription. This switch is assumed to be a stochastic process that facilitates the emergence of ribosomal bimodality.

By using this theoretical framework, we analyzed the experimental data, and found indications of active rRNA degradation being involved in the process of ribosome decrease. First, we found that a strictly non-zero active degradation rate was needed to fit the 16S rRNA-FISH fluorescence intensities over time (Extended Data Fig. 2c; see Supplementary note 2.2). Second, we observed that the degree of ribosomal reduction was ~ 5-fold higher than expected from a dilution-based process by comparing to the experimentally measured optical densities (Extended Data Fig. 2d; see Supplementary Note 3), supporting the necessity of an active degradation mechanism.

According to our analysis, the emergence of bimodal ribosomal distribution is influenced by multiple factors, such as the magnitude of experimental noise (see Supplementary Note 1.5) and the interplay between two key parameters: the rRNA transcription switching rate (*γ_rib_*) and the degradation rate (λ_*a*_) (Extended Data Fig. 2e, see Supplementary Notes 2.1 and 2.2). Numerical analysis of the parameter space suggested that the estimated parameters for *S*.Tm (*γ_rib_* ≈ 1.4*μ*_0_, λ_*a*_ ≈ 3.8*μ*_0_) allow an observable ribosomal bimodality, whereas the parameters estimated for *E. coli* (*γ_rib_* ≈ 3*μ*_0_,λ_*a*_ ≈ 4.2*μ*_0_) do not (Extended Data Fig. 2f; see Supplementary Note 4). In conclusion, our results suggest that the relationship between these two parameters - rRNA transcription switching rate and degradation rate - is critical in driving the emergence of rRNA bimodality in *S*.Tm.

Given the intracellular lifestyle of *S*.Tm during infection, we reasoned that the switch from nutrient surplus to scarcity occurring at the transition phase could mirror the intracellular conditions that *S*.Tm faces during macrophage infection, and hypothesized that it would elicit a similar ribosomal abundance switch. To investigate this hypothesis, we infected J77A-1 macrophages with exponentially growing or stationary phase *S*.Tm. In stationary phase, bacteria displayed 16S^*low*^ rRNA levels (Fig. 1b, OD_600_ = 5). In accordance, during infection with stationary phase *S*.Tm, we did not measure any change in rRNA levels of intracellular bacteria, that remained low and constant (Extended Data Fig. 2g). During infection with exponential phase *S*.Tm, which displays 16S^*high*^ rRNA levels (Fig. 1b, OD_600_ = 0.2), we measured increasing numbers of bacteria that switched to 16S^*low*^ ribosomal levels within infected host cells (Fig. 2c,d). Thus, the intracellular host environment also elicits a switch towards a reduction of rRNA in *S*.Tm.

### Carbon limitation-mediated fitness in a subpopulation of cells during nutritional upshifts

Given the difference in ribosomal levels between the 16S^*high*^ and 16S^*low*^ subpopulations, we hypothesized that phenotypic traits could functionally distinguish these two groups. We aimed at comparing the recovery rates of *S*.Tm cells with different ribosomal levels upon restoration of nutrients during nutritional upshifts. To this end, we grew *S*.Tm in two defined minimal media depleted of carbon or phosphate (Fig. 3a), to achieve high or low ribosomal levels, respectively, as previously reported [38]. First, we quantified 16S rRNA-FISH levels in *S*.Tm grown in carbon or phosphate limitation (Fig. 3b). While in phosphate limitation the majority of cells were 16S^*low*^, in carbon limitation a subpopulation remained 16S^*high*^ (16.3%), displaying a ~ 9-fold increase in rRNA, and resembling the bimodality observed during the transition phase in rich medium (Fig. 1d), although with a smaller fold change between 16S^*high*^ and 16S^*low*^. Second, to test whether differences in rRNA levels correlate with active translation, we examined protein synthesis capacity of *S*.Tm in each medium. Cells were pulsed with the noncanonical amino acid L-Azidohomoalanine (AHA) and protein synthesis was measured using Bioorthogonal Noncanonical Amino Acid Tagging (BONCAT) [39] (Extended Data Fig. 3a,c). Comparing carbon- with phosphate-limited cells, we measured a significantly higher protein synthesis rate in carbon-limited cells (Fig. 3c), suggesting that the 16S^*high*^ subpopulation is translating at a higher level.

**Fig. 3:**
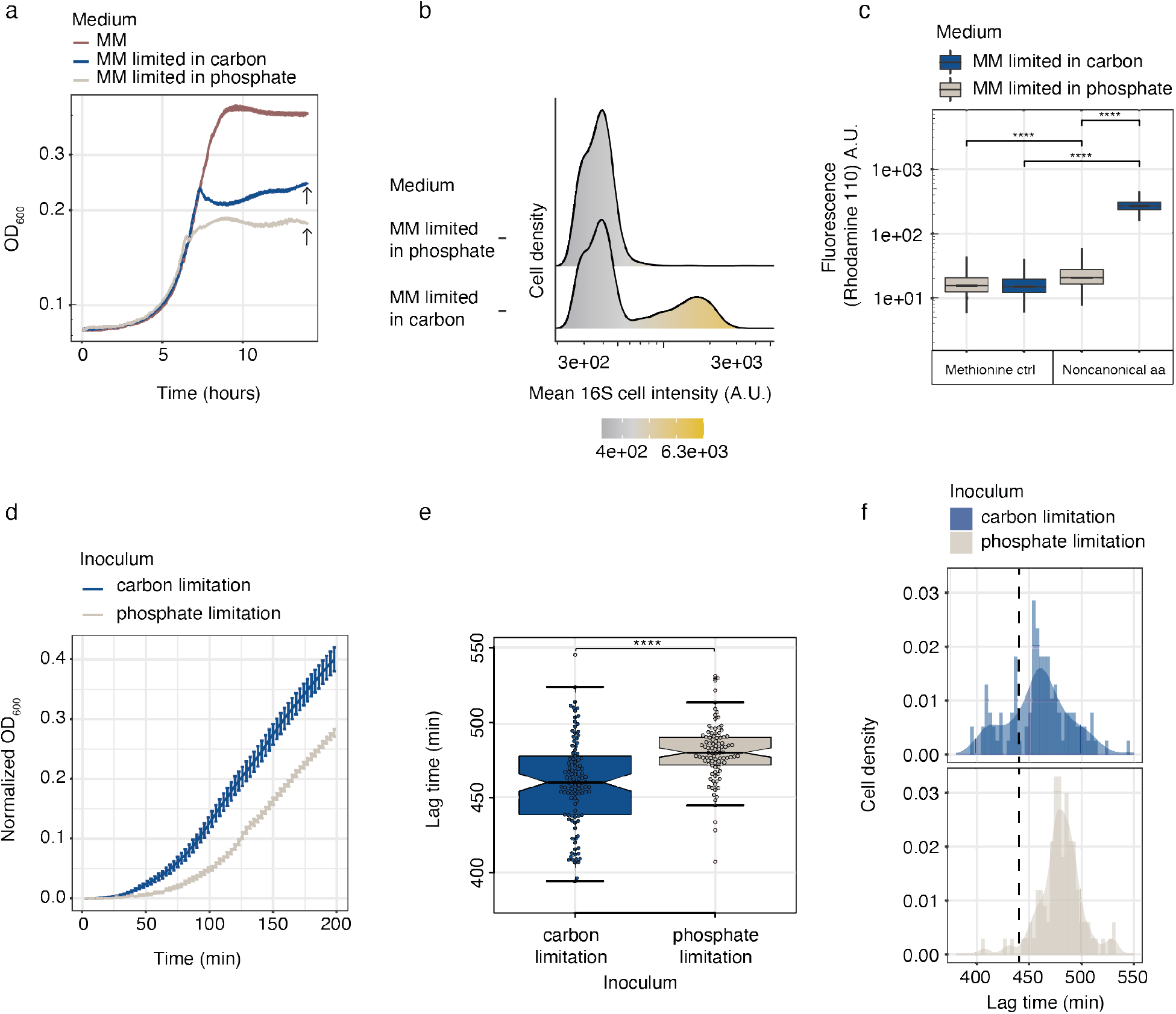
Carbon limitation-mediated fitness in a subpopulation of cells during nutritional upshifts. **a**, Growth curves of *S*.Tm grown in complete minimal medium (MM) (in brown), or modified MM depleted in carbon (in blue) or phosphate (in grey). Arrows indicate the time of harvest for Fig. 3b. **b**, Carbon-limited, but not phosphate-limited bacteria display a bimodal distribution of 16S rRNA. Density histograms show 16S rRNA-FISH mean fluorescence intensity of individual *S*.Tm cells measured from microscopy images from the indicated samples (n = 2212 – 3543 cells). **c**, *S*.Tm translation is lower under phosphate-limitation (in blue) compared to carbon limitation (in grey), as shown by BONCAT analysis (n = 20,356 – 42,613 cells). Methionine controls are shown as background reference. Unpaired t-test: ****, *P* <0.0001. **d**, *S*.Tm cultures grown under carbon limitation (in blue) or under phosphate limitation (in grey) were inoculated into fresh LB medium at the time indicated by the arrows in 3a. Carbon-limited *S*.Tm recover faster than phosphate-limited, as shown by bulk measurements of OD_600_. The absorbance OD_600_ at time 0 was subtracted from each sample to center the data at zero. Error bars indicate means ± SEM of twenty four replicates. **e**, Individual *S*.Tm cells inoculated from carbon-limited MM (in blue) display shorter lag time during nutritional upshifts compared to phosphate-limited (in grey). Nutritional upshifts curves were obtained from individual bacteria sorted into fresh LB medium supplemented with glucose, and used to calculate the individual lag times, shown as boxplots (n = 96 – 114). Unpaired t-test: ****, *P* <0.0001. **f**, Lag times from 3e, presented as density histograms. The first quartile of the carbon-limited sample is indicated with a dashed line.

Next, we tested whether higher rRNA levels matched a specific phenotypic trait, *i.e.*, a growth advantage upon nutritional upshifts. At the indicated time point (see arrows in Fig. 3a), we transferred *S*.Tm cells from depleted media to fresh LB medium supplemented with glucose and measured their growth. From the growth curves, we calculated the lag times and found that carbon-limited cells were able to resume growth significantly faster than phosphate-limited cells (Fig. 3d and Extended Data Fig. 3d). We then repeated the experiment, moving from bulk to single cell measurements, by sorting individual cells into multi-well plates containing fresh LB medium supplemented with glucose, from which single cell-derived growth curves were recorded. In agreement with the bulk measurements, we measured a significantly faster resumption of growth in cells previously subjected to carbon limitation compared to phosphate limitation, both in single cell-derived growth curves (Fig. 3e and Extended Data Fig. 3e) and single cell-derived drop spotting measurements (Extended Data Fig. 3f). In addition, we measured a wider distribution of lag times for carbon-limited cells compared to phosphate-limited cells (*s*^2^ = 1384.2 in carbon, *s*^2^ = 389.5 in phosphate) (Fig. 3f). Interestingly, this variance was driven primarily by a subset of carbon-limited cells that were able to resume growth faster than phosphate-limited cells (cells left to the dashed line, Fig. 3f). Strikingly, the fraction of fast-recovery carbon-limited cells (25.4%) was comparable to the proportion of 16S^*high*^ cells (Fig. 3b), suggesting that these two traits are not independent.

Thus, these results suggest that carbon limitation induces a phenotypic variability in the number of ribosomes that confers an advantage during nutritional upshifts by shortening the lag times in a subpopulation of cells, possibly bearing a higher number of ribosomes compared to the rest of the population.

### DksA-mediated transcriptional shutdown and RNase I-mediated rRNA degradation as drivers of rRNA phenotypic variability

We have shown by theoretical and data analysis (see Supplementary Note 3) a role for active degradation in the formation of the 16S^*low*^ subpopulation, thus, we sought to experimentally identify the genetic factors driving the separation of bacteria into 16S^*high*^ and 16S^*low*^. To approach this in an unbiased manner, we generated a saturated genome-wide library of *S*.Tm mutants via random Tn5 transposition. We grew the transposon mutant library in LB medium to the transition phase, stained for 16S rRNA-FISH and analyzed by flow cytometry [31] (Fig. 4a). 16S^*high*^ and 16S^*low*^ subpopulations were separately sorted and analyzed by Transposon insertion site sequencing (TIS) [40]. By mapping transposon insertion sites, we identified several disrupted genes enriched in the 16S^*high*^ vs. 16S^*low*^ subpopulation (Fig. 4b and Table S2). We focused on the 16S^*high*^ subpopulation, reasoning that this fraction would contain the cells impaired in the genes involved in rRNA reduction. Among the hits, we found several members of the *tol* and *rfa* operons, involved in outer membrane integrity [41] and LPS core biosynthesis [42], respectively. We reasoned that these genes might affect growth rather than be actively involved in rRNA regulation. Two putative candidates, *dksA* and *rna*, were previously suggested to be involved in regulation of rRNA. DksA plays a crucial role in rRNA synthesis during the stringent response by potentiating RNAP response to (p)ppGpp and nucleotide triphosphate (NTP) [24]. *rna* encodes RNase I, an unspecific endonuclease located primarily in the periplasm, with a cytoplasmic form called RNase I* of unknown function [43]. To validate the implication of these candidates in the regulation of rRNA levels, we generated *dksA* and *rna* single-gene knockouts (KOs) and dCas9-mediated knock-downs (KDs) via CRISPR interference (CRISPRi) [44]. In all KO and KD mutants, genetic perturbation of *dksA* or *rna* completely prevented ribosomal reduction and formation of the 16S^*low*^ subpopulation during the transition phase (Fig. 4c,d).

**Fig. 4:**
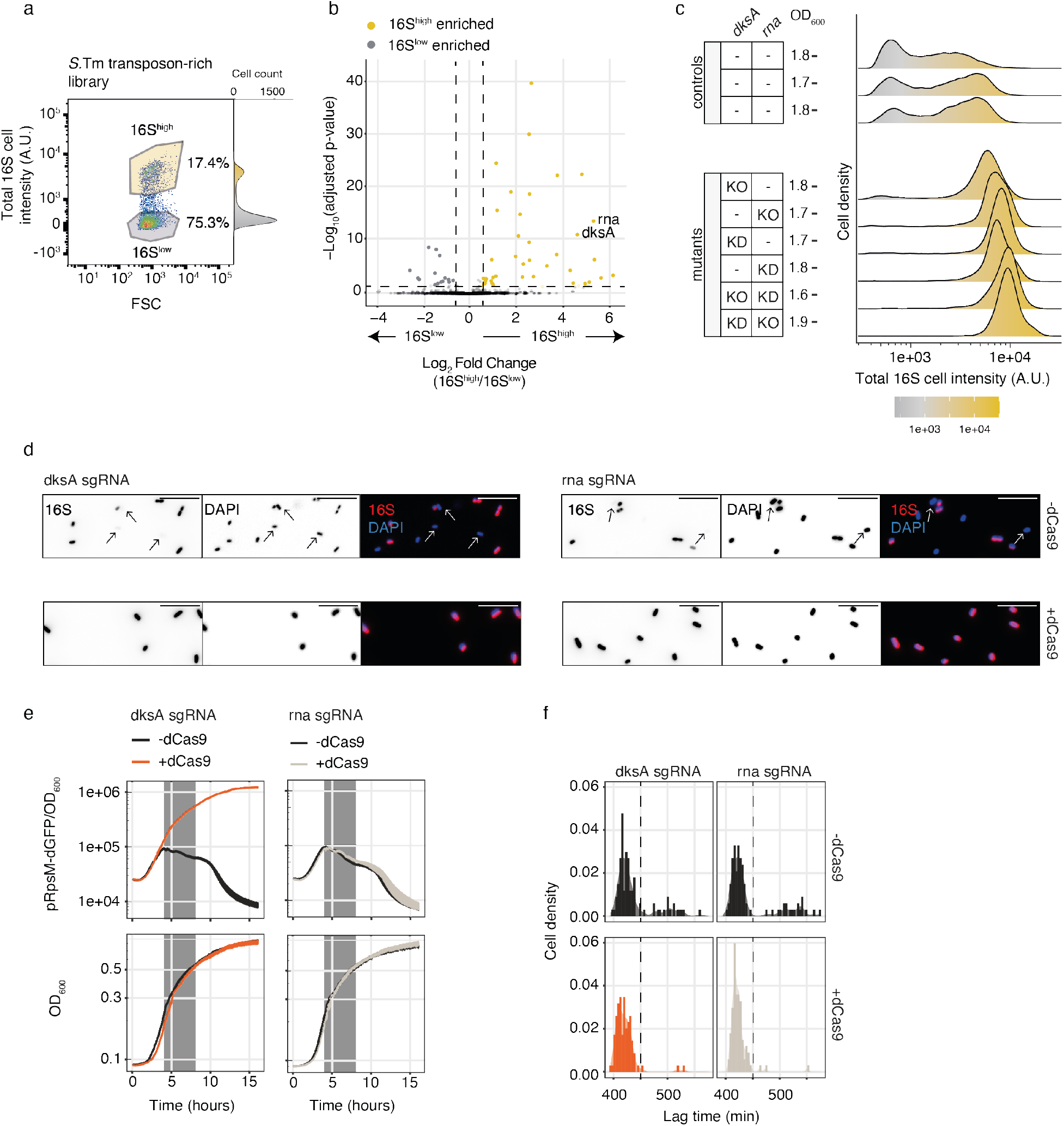
DksA-mediated transcriptional shutdown and RNase I-mediated rRNA degradation as drivers of rRNA phenotypic variability. **a**, *S*.Tm transposon-rich library (~ 250,000 mutants) was grown to the transition phase and analyzed for 16S rRNA-FISH fluorescence by flow cytometry. The plot indicates the gates used for sorting 16S^*low*^ and 16S^*high*^ subpopulations based on 16S rRNA-FISH total fluorescence intensity (per cell) and forward scatter (FSC) areas. Histogram of the data in the y axis is provided. **b**, Volcano plot of mutants enriched in 16S^*low*^ (in grey) and in 16S^*high*^ (in yellow) subpopulations, obtained by TIS analysis. **c**, Bimodality in 16S rRNA signal does not emerge upon genetic perturbations of *dksA* or *rna*. Density histograms show 16S rRNA-FISH total fluorescence intensity (per cell) measured by flow cytometry of individual *S*.Tm cells harvested during the transition phase (n = 10,000 cells). Genetic perturbation of *dksA* or *rna* genes by KO or KD is indicated on the left. Controls are shown in the upper panel. Each sample OD_600_ is indicated on the left. **d**, Representative microscopy images of *dksA* (left side) and *rna* (right side) knocked-down samples (lower row), and the corresponding controls (-dCas9; upper row) from Fig.4c are shown. 16S rRNA-FISH, red; DAPI, blue. Single channels are shown in black. Scale bar, 10 *μ*m. **e**, Transcription of rRNA shuts down at the transition between exponential and stationary phase. pRpsM-dGFP reporter fluorescence was measured during 16 hours of growth in LB and normalized to the OD_600_. Shown are the pRpsM-dGFP fluorescence measurements of *dksA* (in red, left panel) and *rna* (in grey, right panel) knocked-down samples, and the corresponding controls (-dCas9; in black). Error bars indicate means ± SEM of four replicates. The corresponding growth curves measured as OD_600_ are shown in the lower panel. The grey area corresponds to the transition from exponential to stationary phase. **f**, *dksA* (in red, n = 94) and *rna* (in grey, n = 91) knocked-down samples, and the corresponding controls (-dCas9 in black, n = 91, upper row) were grown in LB to the transition phase. Individual bacteria were sorted into fresh LB medium supplemented with glucose to measure nutritional upshifts curves and the corresponding lag times, shown as density histograms.

We further investigated DksA and RNase I mechanisms of action. First, we evaluated whether rRNA transcriptional regulation was disrupted by the absence of DksA. As transcriptional regulation of rRNA and ribosomal proteins is coordinated [45], we constructed a transcriptional reporter where a destabilized GFP (dGFP) was driven by the ribosomal protein RpsM promoter. To allow promoter kinetics measurements, the dGFP was engineered with a C-terminal tail that shortens its half-life to 60 minutes [46]. By analyzing rRNA transcription in an *S*.Tm strain carrying this reporter (pRpsM-dGFP), we measured a steady increase in expression during lag phase, reaching a maximum in 100% of the bacteria during exponential phase. Expression was reduced during the transition phase, dropping to undetectable levels in stationary phase (Extended Data Fig. 4a). To benchmark the kinetics of rRNA transcription using this reporter, we performed single-molecule FISH (smFISH) using a probe specific to the pre-16S rRNA (Extended Data Fig. 4b,c). We validated the kinetics of the fluorescent reporter pRpsM-dGFP, although the dynamics of transcriptional shutdown were delayed compared to the smFISH results, likely due to the half-life of dGFP (Extended Data Fig. 4d). We next introduced the pRpsM-dGFP reporter into *rna*-KD and *dksA*-KD strains and recorded transcriptional activity during their growth. As controls we used the same strains grown in the absence of IPTG, to prevent the KD induction (Fig. 4e, in black). Whereas the control and the *rna*-KD strains showed identical kinetics of rRNA shutdown at the transition phase (Fig. 4e, right panel), rRNA transcription was no longer arrested in the *dksA*-KD (Fig. 4e, left panel). This finding is consistent with previous reports showing higher activity of the P1 promoter of rrnB rRNA operon in a Δ*dksA* background compared to the wild type [24]. To investigate further the specific role of RNase I in rRNA active degradation, we analyzed other *S*.Tm RNases, which did not show any distinct phenotype compared to the wild type (Extended Data Fig. 4e), pointing to the pivotal role of RNase I, likely generating the initial endonucleolytic cleavage necessary for other exonucleases to access rRNA.

We have shown that bimodal rRNA levels are correlated with lag time duration upon nutritional upshifts (Fig. 3f). Due to the absence of the 16S^*low*^ subpopulation in the *dksA*-and rna-depleted strains, we hypothesized that these strains, when subjected to nutritional upshifts, would display a uniform distribution of short lag times. To test this, we grew *dksA*- and rna-KD strains until the transition phase, sorted individual cells, and measured growth recovery. In the absence of IPTG the KD is prevented, thereby, similar to the wild type, we measured a bimodal distribution of cells with short or long lags and large variance (in black, Figs. 4e and 4f; mean = 440.25, SD = 1639.2). Remarkably, KD of *dksA* (in red, Fig. 4f and Extended Data Fig. 4f) or *rna* (in grey, 4f and Extended Data Fig. 4f) resulted in the loss of cells with long lag times, leaving only cells with unimodal short lags and small variance (*mean_dksA_* = 420.28, *SD_dksA_* =371.12; *mean_rna_* = 423.66, *SD_rna_* = 305.2). These results substantiate the dependance between fast growth resumption and 16S^*high*^ status.

In conclusion, we have shown that during the transition to the stationary phase, bacteria switch from 16S^*high*^ to 16S^*low*^ ribosomal levels, by the concerted activity of DksA and RNase I, acting on rRNA transcription and RNA degradation, respectively. Importantly, ribosomal phenotypic variability was correlated with phenotypic fitness during nutritional upshifts, suggesting a bet-hedging behavior, where a subpopulation maintaining higher levels of ribosomes will be favored upon nutrient restoration.

## Discussion

Bacteria constantly experience changes in the environment, with sudden nutrient fluctuations, to which they rapidly need to adapt, *e.g.*, by adjusting ribosomal levels to optimize growth rates. In this study, we investigate how *S*.Tm ribosomal levels change outside of the steady state. Our results demonstrate that in the transition between exponential growth and stationary phase, when bacteria slow their growth, ribosomal levels are reduced by combining transcriptional shut-down and rRNA degradation. Within the population, this phenomenon is not synchronized and gives rise to two subpopulations of *S*.Tm cells harboring different levels or ribosomes (16S^*high*^ and 16S^*low*^), and we suggest that in this ribosomal cell-to-cell heterogeneity underlies a fitness advantage.

By moving from bulk to single cell measurements, our findings expand previous observations on ribosomal decrease in slow-growing bacteria and in non-growing bacteria subjected to various types of starvation [3, 5, 10, 12], and uncover a bimodal behavior of *S*.Tm. In previous work, highly variable levels of ribosomal amounts have been reported in *Pseudomonas Aureoginosa* cells during starvation [47], however not investigated. The bimodality observed in *S*.Tm rRNA levels was reproducible in several other *S*.Tm strains, however not in the Gram-positive *S. aureus*, nor in *E. coli* where the decrease was less abrupt.

Our theoretical framework suggests that bimodality arises from a stochastic switching mechanism during the transition from exponential to stationary, forming the basis for bimodality. Fitting our experimental measurements of rRNA-FISH with the theoretical framework revealed that the switching mechanism only does not support the emergence of bimodality, and that an rRNA degradation process must be included. Our results showed that the emergence of ribosomal bimodality is closely tied to the relationship between a few key parameters, the switching rate *γ_rib_* and the degradation rate *λ_α_*, and suggested that the estimated parameters support observable bimodality in *S*.Tm, but not in *E. coli* (Extended Data Fig. 2f), aligning with the experimental findings.

Notably, although rRNA is reported to be degraded in response to certain physiological conditions [3, 5, 10, 12], the drivers of this process were not conclusively characterized [2]. Our study identified DksA and RNase I as crucial players in this process, acting on rRNA transcriptional shutdown and rRNA degradation, respectively. Importantly, KD and KO of each of these genes impaired the process of ribosomal reduction observed during the transition phase, thus resulting in the loss of bimodality in the population. An important player involved in rRNA regulation, namely in rRNA transcriptional shutdown, is the stringent response, a broadly conserved bacterial stress adaptation mechanism occurring at the onset of stationary phase in response to several starvation signals, in which the alarmone molecule (p)ppGpp is synthesized [48]. Acting in synergy with the stringent response, DksA was previously reported to sensitize RNAP to (p)ppGpp, and assist in stopping rRNA transcription [24]. In our experiments, rRNA transcription failed to shut down in the absence of DksA (Fig. 4e), resulting in continuous synthesis of new ribosomes. Further experiments will be required to identify the upstream regulators of DksA and RNase I. Similarly, future experiments may indicate which isoform of RNase I is involved, the cytosolic or the periplasmic, perhaps becoming active due to increased membrane permeability.

Finally, we demonstrated that phenotypic variability in ribosomal levels confers a fitness advantage. During nutritional upshifts, bacteria behave differently depending on the nutrient limitation to which they were previously subjected to [38]. We showed that compared to phosphate, carbon limitation conditions conferred higher levels of translating ribosomes, and, upon restoration of nutrients, a fitness advantage by shortening recovery times. Importantly, our single cell measurements revealed that only a subpopulation of cells was characterized by reduced lag times (Fig. 3f). Remarkably, the proportion of these cells was similar to the proportion of cells harboring high ribosomal levels, suggesting a correlation between ribosomal content and lag time. We further confirmed this correlation using *dksA* and *rna* KO and KD mutants, exclusively constituted of the 16S^*high*^ subpopulation, as they showed a homogeneous behavior in nutritional upshifts with short recovery times (Fig. 4f).

In summary, we have shown that DksA and RNase I play a role in ribosomal reduction, a conserved process that occurs when bacteria slow their growth in response to nutrients scarcity. We showed that this mechanism is active in only a subpopulation of cells, while another subpopulation maintains high ribosomal levels for a longer period of time, and this fraction of cells is favored during nutritional upshifts. We suggest that the phenotypic variability in ribosomal levels follows a bet-hedging strategy, where a subpopulation of bacteria will be able to resume growth faster upon restoration of nutrients. Future studies may implicate the physiological relevance of our findings to adaptation and survival within the host. Inside the gut, enteric bacteria go through daily feast and famine cycles, for which tight regulation of ribosomal amounts could contribute to better survival. Maintaining phenotypic heterogeneity at the population level might be a strategy to be prepared for different scenarios, *i.e.*, feast or famine. Furthermore, being capable of utilizing the building blocks made available by ribosomal degradation could particularly benefit intracellular pathogens such as *Salmonella*, given their intracellular life cycle during infection, where following host invasion they reside within a modified phagosome with limited access to nutrients.

## Methods

### Bacterial strains and plasmids

All strains used in this study are listed in Supplementary Table 1. *E. coli* K-12, *S. aureus, S*.Tm S4/74, *S*.Tm D23580, *S*.Tm ATCC 14028s wild type (WT), and the *S*.Tm Single Gene Deletion (SGD) mutants [49] Δ*rnb*, Δ*rph*, Δ*vacB*, and Δ*yfgH* were used. *S*.Tm Δ*dskA* and Δ*rna* mutants were constructed using λ-red recombinase gene replacement method [50] using WT ATCC 14028s strain as background. Deletion was verified by PCR using primers listed in Supplementary Table 1, using gene-specific primers as forward, and Kan_KO or Cam_KO as reverse according to the resistance carried (kanamycin or chloramphenicol). *S*.Tm knock-down (KD) strains were constructed by insertion of a Mobile-CRISPRi module as described below.

### Construction of the Mobile-CRISPRi vectors

sgRNAs were designed as previously described using CHOPCHOP [51] (https://chopchop.cbu.uib.no/). sgRNAs were cloned into the BsaI sites of pJMP1339 (addgene #119271) by ligating pairs of annealed oligos (Supplementary Table 1). Oligos included 5’ overhangs complementary to the sticky ends generated by BsaI-HF (NEB, R3733S) restriction enzyme. Pairs of oligos were phosphorylated at 37°C for 30 min in 10 *μ*l final volume containing 10 *μ*M of each oligo, 1x T4 DNA ligation buffer (NEB, B0202S), and 5U T4 PNK (NEB, M0201L). Then, they were denatured at 95°C for 5 minutes and annealed by ramping down the temperature to 25°C, decreasing it 5°C/min. Two *μ*l of a 1:250 dilution of the annealed oligos were ligated to 50 ng of dephosphorylated BsaI-digested vector at 16°C overnight in 20 *μ*l final volume containing 1x T4 DNA ligation buffer (NEB, B0202S), and 400U T4 DNA ligase (NEB, M0202L). Ligations were purified and transformed by electroporation in a MFDpir24 strain [52], a pir+ strain diaminopimelic acid (DAP)-auxotroph containing the RP4 transfer machinery (kind gift of Prof. Jean-Marc Ghigo, Pasteur Institute), using 2 mm electroporation cuvettes (Cell Projects, EP-102) and electroporation program Ec2 (V = 2.5 kV) with MicroPulser Electroporator (Biorad). Plasmid sequences were confirmed by sanger sequencing.

### Construction of the Mobile-CRISPRi strains and mating assays

Tn7-based Mobile-CRISPRi strains were constructed by triparental mating as previously described [53], with few modifications. All Tn7 matings used MFDpir24 strain, transformed with either a Tn7 transposase plasmid pJMP1039 (addgene, #119239) or a transposon plasmid (derivatives of pJMP1339, addgene #119271, containing sgRNAs). The so-obtained MFDpir24 donor strains (transposon and transposase mating donors), were grown at 37°C overnight in LB supplemented with 300 *μ*M DAP (Sigma, D1377) and 100 *μ*g/ml ampicillin. The recipient strain *S*.Tm was also grown in LB at 37°C for ~ 16 h. Of each donor and recipient strain, 100 *μ*l was added to 700 *μ*l LB and mixed by pipetting. Mixes of donor and recipient strains were pelleted for 2 min at 7,000*xg*, washed twice with 1 ml LB, resuspended in 30 *μ*l LB, spotted on a LB plate, and incubated at 30°C overnight. Spots were then resuspended in 100 *μ*l PBS and spread onto LB agar plates that selected for the Mobile-CRISPRi plasmid and recipient (LB + 30 *μ*g kanamycin) without DAP. Insertion of the module was verified by PCR using primers listed in Supplementary Table 1.

### Construction of fluorescent reporter vectors

Plasmid pFPV25.1_dGFPmut3-AAV (addgene, #187377) (referred to as pRpsM-dGFP) was cloned by Gibson assembly using pFPV25.1 (addgene, #20668) as backbone. pFPV25.1 was digested with XbaI-HF (NEB, R0145S) and HindIII-HF (NEB, R3104S) restriction enzymes. The insert was amplified using pFPV25.1 as template with primers listed in Supplementary Table 1. Plasmid sequence was confirmed by sanger sequencing and next-generation sequencing (NGS).

### Growth media and conditions

Unless otherwise mentioned, bacterial strains were cultured for 16 hours at 37°C in LB medium, diluted 1:1,000 and grown in LB medium. For minimal media (MM) experiments, cells were grown in modified MOPS media (9.5 mM NH_4_Cl, 50 mM NaCl, 0.5 mM CaCl_2_, 0.525 mM MgCl_2_, 0.276 mM K_2_SO_4_), glucose (0.4%, Sigma G8270), and phosphate (1.32 mM K_2_HPO_4_, Sigma P3786) added separately. For carbon and phosphate limiting media, glucose and K_2_HPO_4_ concentrations were reduced by 5-fold and 10-fold (0.08% and 0.132 mM, respectively). For the comparison or rRNA-FISH measurements and *r* ratios from literature [4], *E. coli* was grown in minimal medium based on Miller’s M6350 or in rich defined medium (RDM), a MOPS buffered medium supplemented with micronutrients, amino acids, and vitamins, described by Neidhardt et al [54].

### CFUs

CFUs were quantified by serial dilution on LB plates containing the appropriate antibiotics and counted with Scan 500 automatic colony counter (Interscience).

### FISH

FISH probes used are listed in Supplementary Table S1. 16S, pre-16S, and 23S rRNA probes were synthesized with a 5’ fluorophore label (IDT). FISH was performed as described previously [32], with few modifications. Briefly, cells were fixed in 3.7% PFA (ThermoFisher, 28906), and permeabilized in 70% ethanol overnight at 4°C. Probes were hybridized overnight at 37°C in hybridization buffer containing 10% dextran sulphate (Sigma, D8906-5G), 25% formamide (Roche, 11814320001), 1 mg/ml *E. coli* tRNA (Sigma, R4251), 2x SSC (Ambion, AM9765), 0.02% BSA (Thermo Fisher, AM2616), 2 mM vanadyl ribonucleoside complex (NEB, S1402S). The cells were then washed 4 times in wash buffer containing 25% formamide (Roche, 11814320001), 2x SSC (Ambion, AM9765). In the last wash, 10 *μ*g/ml DAPI (Roche, 10236276001) was supplemented. Cells were mounted on microscopy slides using GLOX anti-fade buffer containing 10 mM Tris pH 8 (Thermo Fisher, AM9856), 2x SSC (Ambion, AM9765), 0.4% glucose (Merk, 50-99-7), 37 mg/ml glucose oxidase (Sigma, G2133), and 100 mg/ml catalase (Sigma, C3515). Imaging was performed as described below, on the same day as the mounting.

### Microscopy

For determining the spatial localization of 16S and pre-16S rRNAs in the bacterial cell (Fig. S1c), imaging was performed with a CSU-10 spinning disk system (Yokogawa Electric Corp., Tokyo, Japan) assembled on an Eclipse Ti-E inverted microscope (Nikon), equipped with a x100 NA 1.45 oil-immersion objective, laser set (405, 561, 638 nm), and a pinhole of 50 mm. The rest of the microscopy experiments were performed on an inverted epifluorescence microscope (Eclipse Ti2-E, Nikon), equipped with a x100 NA 1.45 oil-immersion objective, filter sets (DAPI, GFP, Cy3, Cy5/Cy5.5), and either an EMCCD camera (iXon Ultra 888, Andor) or a CMOS camera (Prime 95C, Photometrics). Exposure times above 300 ms were avoided to minimize photobleaching. For segmentation purposes, the DIC channel was also acquired. For the in vitro experiments, image stacks consisting of 7-9 z focal planes with 200 nm spacing were acquired for several xy slide positions per sample (typically 10-20 positions). For the infection experiments, image stacks consisting of 17 z focal planes with 300 nm spacing were acquired for several xy slide positions per sample (typically 30 positions).

### Image analysis

Bacterial cells were segmented using the machine-learning software Ilastik [55], following the “pixel and object” classification workflow. The ilastik binary masks were further processed with a custom matlab script aimed at removing spurious objects (small and big objects), and at curating the segmentation of high-density regions via a watershed algorithm. To quantify cell fluorescence, we used a custom matlab script measuring average cell fluorescence, subtracted of the background (fluorescence outside of cells). For pre-16S rRNA quantification, we used the MATLAB-based spot-recognition software Spätzcells [32]. Briefly, in each z focal plane of an image stack, the local maxima are identified above a user-defined threshold based on the negative control (unstained control). Local maxima from different z planes that correspond to the same spot are joined together, and the plane with the highest intensity is defined as the in-focus plane. In this plane, the fluorescence intensity profile of each spot is fit to a 2D Gaussian to estimate position, area, peak height, and spot intensity. Spots are assigned to cells using the segmentation masks. False positive spots are discarded using the negative control sample (unstained control), setting a cutoff so that only spots brighter than the 99% of the spots detected in the negative control are considered true positives. Fluorescence corresponding to single RNA molecules is calculated using as reference the sample with the lowest transcriptional level (OD_600_ = 1.8).

### Propidium Iodide staining

Bacteria were incubated in a 2x SSC solution containing 500 nM propidium iodide (Thermo Fisher, P3566) for 5 minutes at RT, before acquisition on FACS. As positive control (dead cells), bacteria were heated at 95°C for 5 minutes.

### Flow cytometry

Fluorescence acquisition was performed on a BD FACSAria III (BD Biosciences) using a 70 *μ*M nozzle, or on a Amnis ImageStream (Luminex). Sorting was performed on a BD FACSAria III (BD Bioscience), using single cell mode. Flow cytometry data was analyzed using FlowJo Software (BD).

### RT-qPCR

One million cells per sample were sorted on a BD FACSAria III sorter, and pelleted in 3% BSA. De-crosslinking was performed at 37°C for 30 minutes via proteinase K digestion in 50 *μ*l of proteinase K buffer (Tris 100 mM, EDTA 10 mM, NaCl 50 mM, 7M urea) containing 1.2 mg/ml of proteinase K (Thermo Fisher, AM2546). To this reaction, 650 *μ*l of Trizol (Thermo Fisher, 15596026) was added, bacteria lysed with Omnilyse (ClaremontBio, 01.340.48), and RNA extracted with miRNAsy kit (Qiagen, 217004) following manufacturer’s instructions (including the optional DNAse digestion on column). A second DNA digestion was performed using Turbo DNAse (Thermo Fisher, AM2238) following manufacturer’s instructions. cDNA was generated using SuperScript III (Thermo Fisher, 18080093) and random hexamers according to manufacturer’s instructions. RT-qPCR was performed using Fast SYBR Green Master Mix (Thermo Fisher, 4385616) on a QuantStudio 5 instrument (Applied biosystems).

### RT-qPCR

J77A-1 (8×10^5^ cells/well) were plated in untreated six–well plates (Greiner, 657185) supplemented with DMEM (Thermo Fisher, 41965-039) containing 10% FBS (Biological Industries, 04-007-1A) and 1 mM sodium pyruvate (Biological Industries, 03-042-1B). LB-grown cultures of *S*.Tm were washed with 2x PBS before infection. J77A-1 were infected at a multiplicity of infection (MOI) of 3:1 and spun down for 5 min at 400xg to synchronize internalization. After 30 minutes, cells were washed with medium containing 50 *μ*g/ml gentamicin to remove not internalized *S*.Tm. Fresh medium containing 50 *μ*g/ml gentamicin was then added back to the cells for the duration of infection.

### BONCAT

For BONCAT experiments, cells were grown in modified MOPS media (9.5 mM NH_4_Cl, 50 mM NaCl, 0.5 mM CaCl_2_, 0.525 mM MgCl_2_, 0.276 mM K_2_SO_4_), glucose (0.4%, Sigma G8270), and phosphate (1.32 mM K_2_HPO_4_, Sigma P3786) added separately. For carbon and phosphate limiting media, glucose and K_2_HPO_4_ concentrations were reduced by 5-fold and 10-fold (0.08% and 0.132 mM, respectively). Bacteria were pulse-labeled for 60 min by supplementing 1 mM L-AHA (Click Chemistry Tools, 1066-100) or L-methionine (Sigma, M9625-5G) as control, as previously described [56]. Briefly, cells were then fixed in 3.7% PFA for 2 hours at 4°C, and washed 3x in PBS to remove excess L-AHA. Dehydration and permeabilization was performed by three step-wise incubations in increasing concentration of ethanol (50%, 80%, 96%), each incubated at room temperature for 3 minutes. Click reaction was performed at room temperature for 30 minutes in 250 *μ*l of PBS containing 5 mM sodium ascorbate (sigma, A4034-100G), 5 mM aminoguanidine hydrochloride (sigma, 396494-25G), 100 *μ*M CuSO_4_ (sigma, 451657-10G), 0.5 mM THPTA (sigma, 762342-100MG), and 24 *μ*M carboxyrhodamine 110 Alkyne (Click Chemistry Tools, TA106-1). Cells were washed 3x in PBS before acquisition via FACS or ImageStream.

### Nutrient upshifts growth measurement

For bulk measurements, carbon- or phosphate-limited cells were mixed with 4x volumes of fresh prewarmed LB medium supplemented with 0.4% glucose and placed in 96-well microplates (Invitrogen, M33089). For single cell measurements, individual cells under carbon- or phosphate-limitation treatments were directly sorted using “single cell” purity mode in a well of a 384-well microplate containing fresh pre-warmed LB medium supplemented with 0.4% glucose. For both bulk and single cell measurements, cells were grown with agitation at 37°C in an imaging multimode reader (Cytation 5, Agilent). Cell growth was monitored every 5 min by measuring absorbance at 600 nm. Lag times were extracted from the growth curves using the R package growthrates [57].

### Drop spotting for nutrient upshifts

Bacteria from single cell nutrient upshifts described above were spotted on LB agar plates 3 hours after being placed in fresh prewarmed LB medium supplemented with 0.4% glucose. Plates were incubated at 30°C until colonies were visible.

### Transposon library construction

Transposon library of *Salmonella* was generated using EZ-Tn5 <KAN-2>Tnp Transposome kit (Lucigen, TSM99K2) following manufacturer’s instructions. Briefly, 300 ml of *S*.Tm was grown in LB at 37°C to a final OD_600_ of 0.8. Cells were washed twice in 180 ml of ice-cold wash solution (300 mM sucrose), and then resuspended in 1 ml of wash solution. Each 100 *μ*l aliquot was transformed with 0.1 pmol of EZ-Tn5 <KAN-2>Tnp Transposome using 2 mm electroporation cuvettes (Cell Projects, EP-102) and electroporation program Ec2 (V = 2.5 kV) with MicroPulser Electroporator (Biorad). Each reaction was allowed to recover in 500 *μ*l SOC (2% tryptone, 0.5% yeast extract, 10 mM NaCl, 2.5 mM KCl, 10 mM MgCl_2_, 10 mM MgSO_4_, 20 mM glucose) for 45 minutes at 37°C, and then plated in a 15 cm^2^ LB agar plate supplemented with kanamycin 30 *μ*g/ml. On the following day transformants were scraped off of the LB plates and pooled into a library of approximately 250,000 transformants.

### TIS sample preparation and illumina sequencing

DNA was extracted from 5 million sorted bacteria using DNeasy Blood & Tissue (Qiagen, 69504) following manufacturer’s instructions, with an overnight proteinase K digestion step (which also serves as PFA de-crosslinker), and resuspension in 200 *μ*l of resuspension buffer. DNA ( 20 ng per sample) was processed as previously described [58], with few modifications. Briefly, DNA was sonicated at 4°C (program: 6 cycles of 30 s on, 60 s off) on a Bioruptor Plus (Diagenode), achieving a size distribution between 200-1000 nucleotides (nt), with 450 nt average size. The sonicated DNA was end-repaired for 30 minutes at 20°C in 200 *μ*l final volume containing 1x end-repair buffer and 10 *μ*l of end-repair mix (NEB, E6050L). The reaction was cleaned in 2x AmpureXP Beads (Beckman Coulter) and eluted in 37.5 *μ*l. A-tailing was performed 20 min at 37°C in 50 *μ*l final volume, containing 1x NEBuffer 2 (NEB, B7002S), 1 mM ATP, and 125U of Klenow fragment (NEB, M0212M). The reaction was cleaned in 2x AmpureXP Beads (Beckman Coulter) and eluted in 44 *μ*l. DNA was then dephosphorylated overnight at 37°C in 50 *μ*l final volume, containing 1x rCutSmart buffer (NEB, B6004S), and 5U of quick CIP (NEB, M0525L). The day after, the reaction was heat-inactivated for 2 minutes at 80°C, cleaned in 2x AmpureXP Beads (Beckman Coulter), and eluted in 18 *μ*l. The DNA template was ligated for 20 min at 25C in 50 *μ*l final volume, containing 0.22 *μ*M Illumina-compatible forked indexed adapters, 1x buffer, and 5 *μ*l of quick ligase (NEB M2200L). To release the non-ligated strand of the adaptor, DNA was denatured at 96°C for 5 min, cleaned with 0.8x AmpureXP beads (Beckman Coulter), and eluted in 30 *μ*l. Two PCR reactions were used for TIS library construction. The first PCR reaction was performed in triplicate in 25 *μ*l final volume containing 0.5 *μ*M TN bait primer 1 (Supplementary Table S1), 0.5 *μ*M Illumina enrichment primer, 1x KAPA HiFi HotStart ReadyMix (Roche, KK2601) and 10 *μ*l DNA template. PCR program: 60s 98°C, 15 cycles of 15s 98°C, 20s 68°C, 60s 72°C, and final extension of 60s 72°C. Following the first PCR reaction, the amplified DNA from the three reactions was pooled and cleaned with 1x AmpureXP beads (Beckman Coulter), and the eluate was used as a template in the second PCR reaction. Conditions for the second PCR reaction were identical to the first, with the following differences: reaction was performed in a single 50 *μ*l reaction and TN bait primer mix 2 consisted in a mix of staggered nested primers (Supplementary Table S1). Libraries were quantified with NEBNext Library Quant Kit for Illumina (NEB, E7630L) and sequenced on a Novaseq machine with the following settings: R1=85, I1=8, and R2=37.

### TIS analysis

Read1 was used to extract reads mapping to the transposon region, by searching for the sequence AGATGTGTATAAGAGACAG (3 mismatches allowed). The region matching to the transposon was trimmed from R1 before mapping, resulting in 37 nucleotides long reads. Transposon-mapping R1 and R2 reads were then mapped to the *S*.Tm 14028S genome with the program STAR, using the following parameters: `-genomeSAindexNbases 10, - outFilterMismatchNmax 3, –outFilterMatchNmin 15`. Multimappers were excluded from the analysis. Read 1 was used to identify the disrupted gene. Read 2, containing the random sonication point, was used as a unique molecular identifier (UMI) to collapse PCR duplicates. On average 10 integrations per gene were recovered in each sample. Mutant enrichment analysis was performed with DESeq2 using UMI counts as input [59].

### GFP fluorescent measurements

Cultures of *S*.Tm carrying pRpsM-dGFP reporter plasmid were grown in LB supplemented with 100 *μ*g/ml ampicillin in 96-well microplates (Invitrogen, M33089) with agitation at 37°C for 16 hours in an imaging multimode reader (Cytation 5, Agilent). Optical density at 600 nm (OD_600_), and GFP fluorescence (488/10 filter set) were measured every 5 min. To obtain individual cell measurements, bacteria were also acquired via FACS.

---

**Extended Data Fig. 1:**
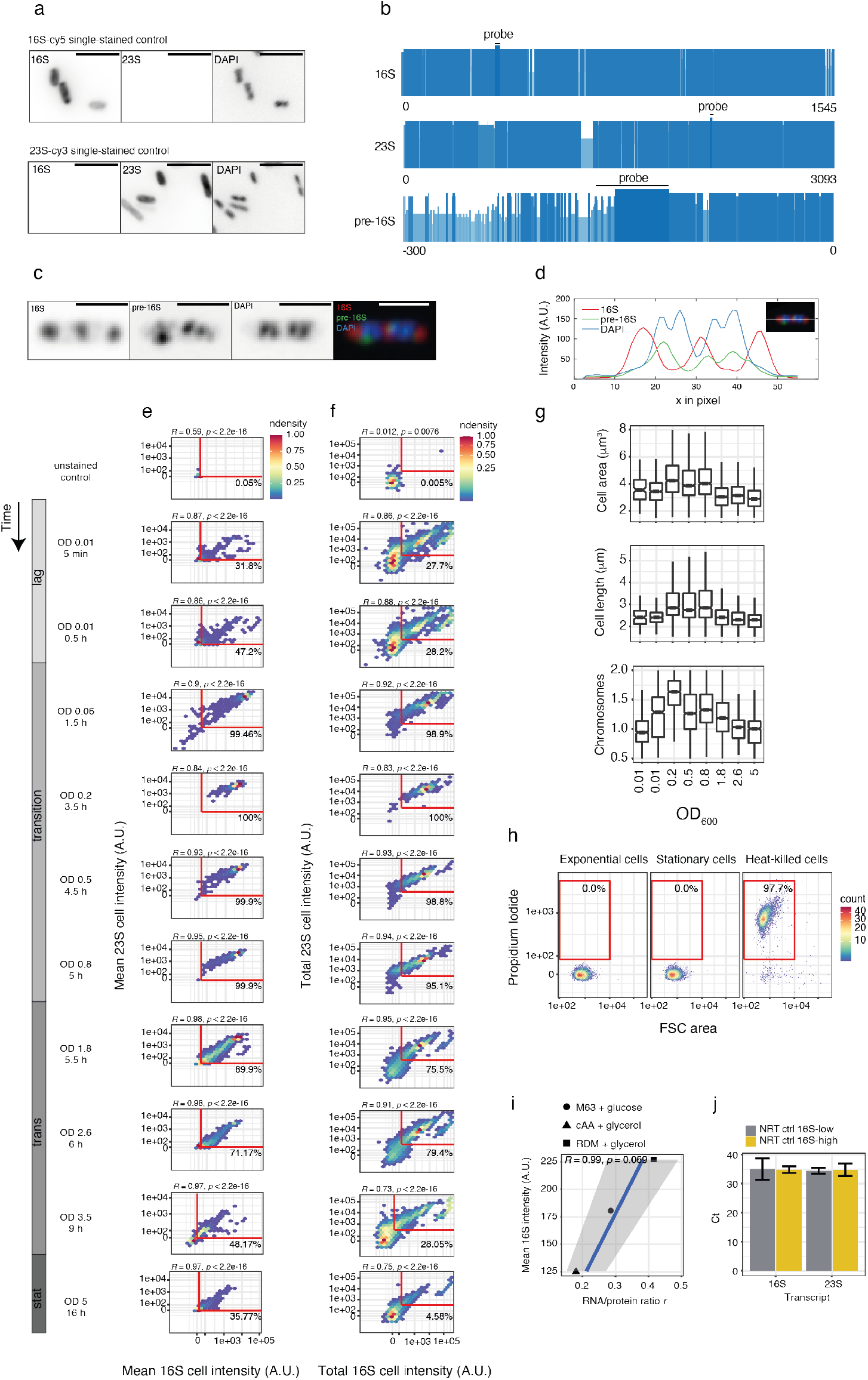
Quality controls of rRNA-FISH. **a**, 16S-Cy5 and 23S-Cy3 dyes are spectrally distinct as shown by the single-stained controls. Single-stained controls for 16S-Cy5 (upper panel) and 23S-Cy3 dyes (lower panel). Cy5, Cy3, and DAPI channels are shown separately in black. Scale bar, 5 *μ*m. **b**, Consensus from multiple sequence alignment of *S*.Tm and *E. coli* 16S, 23S, and pre-16S rRNA operons, shown in blue shades. The 16S, 23S, and pre-16S probes binding sites are shown in black. **c**, 16S rRNA signal localizes to cytosolic regions corresponding to rRNA loaded into ribosomes, as opposed to pre-16S rRNA colocalizing with DAPI and corresponding to newly transcribed rRNA. Representative microscopy image of an exponential phase *S*.Tm bacterium stained with 16S-Cy5 in red, pre-16S-Cy3 in green, and DAPI in blue. Single channels in black. Scale bar, 2.5 *μ*m. **d**, Intensity profile of each fluorescent channel, color-coded as in Extended Data Fig. 1c. The intensity profile was measured across the horizontal axis of the cell as indicated by the white line. **e**, 16S and 23S rRNA signals highly correlate. 2D distributions of *S*.Tm 16S and 23S rRNA mean fluorescence intensities measured from microscopy images from the indicated ODs and time points. Non-stained bacteria (unstained control) were used to gate above background fluorescence (in red, with percentages) (n = 967 – 6,025 cells). **f**, 2D distributions of *S*.Tm 16S and 23S fluorescence intensity areas measured by flow cytometry from the indicated ODs and time points, using DAPI+ gated population. Non-stained bacteria (unstained control) were used to gate above background fluorescence (in red, with percentages) (n = 20,000 cells). **g**, Cellular characteristics of bacteria in different growth phases from Extended Data Fig. 1e. Boxplots show cell area (upper panel), cell length (middle panel), and number of chromosomes (lower panel). To estimate the number of chromosomes, the average DAPI fluorescence was multiplied by the area of the cell and normalized by the median of cells in stationary phase (OD = 5, non-dividing cells with 1 copy of the chromosome). **h**, Cells in stationary phase maintain the membrane intact. Exponential, stationary phase, and dead cells were stained with propidium iodide that can only enter cells with a broken membrane. The gating shows the percentage of propidium iodide-positive cells (red square) (n = 10,000 cells). **i**, 16S-FISH correlates with RNA/protein ratio measurements from literature [1]. *E. coli* exponentially growing at different rates in different media is characterized by different RNA/protein ratios. Bulk averages of 16S rRNA measured by 16S rRNA-FISH and acquired by ImageStream are plotted against RNA/protein ratios, showing a linear correlation. Solid triangle, M63 with glycerol (0.5%, w/v) (growth rate λ = 0.4 hours^-1^); solid circle, M63 + cAA (0.2% casamino acids, w/v) with glucose (0.5%, w/v) (growth rate λ = 1 hours^-1^); solid square, RDM with glycerol (0.5%, w/v) (growth rate λ = 1.31 hours^-1^). **j**, RT-qPCR samples are free of gDNA. No RT (NRT) controls of samples from Fig. 1g are shown for 16S^*low*^ (in grey) and 16S^*high*^ (in yellow) samples, respectively. Values are averaged across three technical replicates. Error bars indicate means ± SEM.

**Extended Data Fig. 2:**
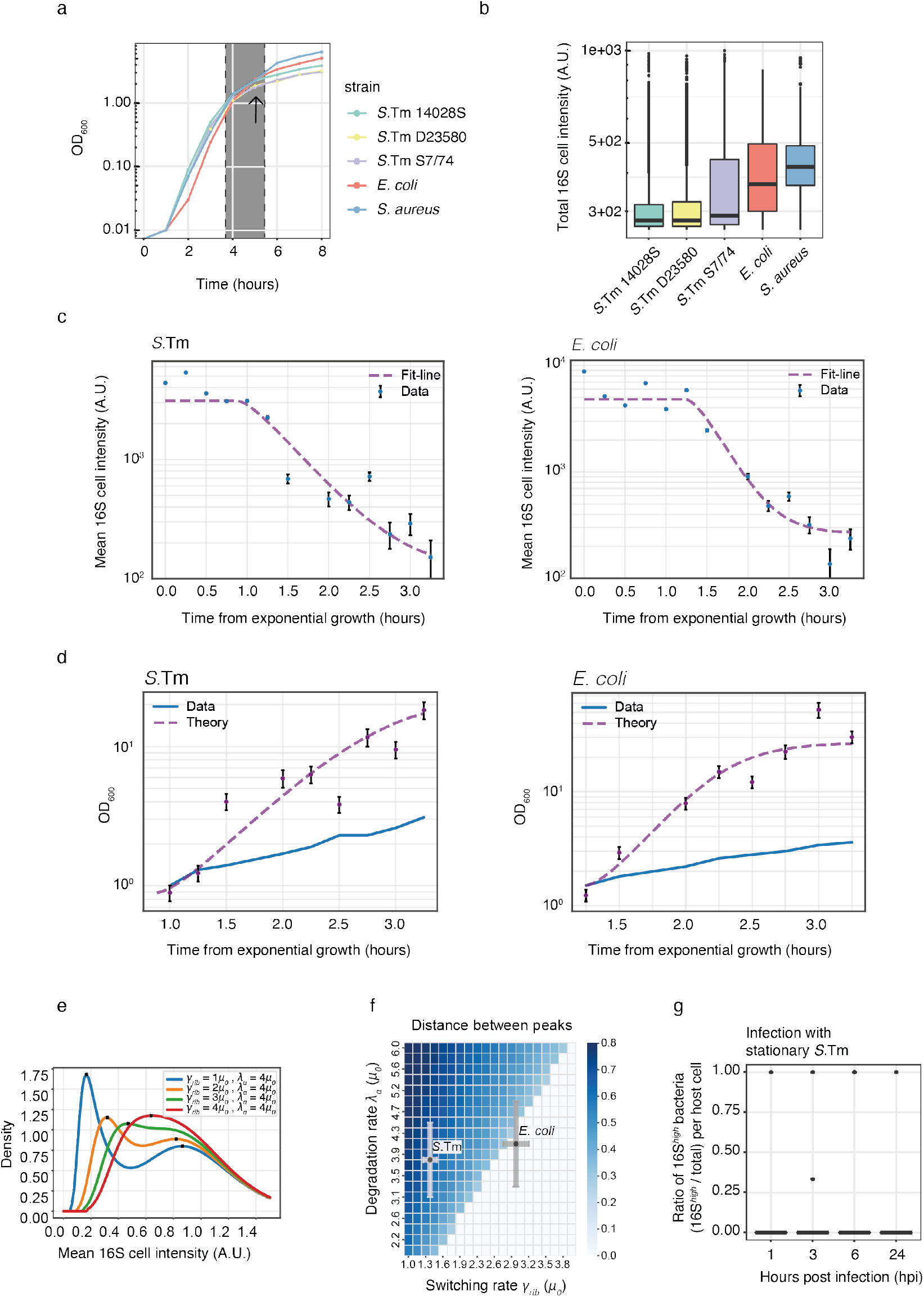
rRNA-FISH in different bacteria and in infection. **a**, Growth curves measured as OD_600_ for the strains shown in Fig. 2a, data points every hour. The grey area corresponds to the transition point before entry into stationary phase, and the arrow indicates the time point assayed in Fig. 2a. **b**, Boxplot showing total 16S rRNA per cell, from the samples as in Fig. 2a. **c**, Infection with stationary phase bacteria. The fraction of 16S^*high*^ bacteria over the total measured per infected J77A-1 macrophage cell is shown. Samples were taken at 1, 3, 6, and 24 hours post infection (n = 27 – 49 host cells, n = 35 – 121 bacteria). **d**, *OD_exp_*(*t*) fitting using eq. 25. A fit of the theoretical expression eq. 25 (dashed purple line) to the measured *OD_exp_*(*t*) (blue circles) for the two bacteria. For *S*.Tm, the fitting parameters came as *γ_rib_* = 1.4*μ*_0_ ± 0.1*μ*_0_ and *t*_0_ = 0.85 ± 0.08hr. For *E. coli*, the fitted parameters came as *γ_rib_* = 3.0*μ*_0_ ± 0.2*μ*_0_ and *t*_0_ = 1.23 ± 0.02hr. **e**, 〈*r*(*t*)〉 fitting using eq. 20. A fit of the theoretical expression eq. 20 (dashed purple line) to the measured mean ribosome concentration (blue circles) for the two bacteria. The y-axis represents the mean ribosome fluorescence concentration in arbitrary units. For *S*.Tm, the fitting parameter came as *λ_a_* = 3.8*μ*_0_ ± 0.7*μ*_0_. For *E. coli*, the fitted parameter is *λ_a_* = 4.2*μ*_0_ ±0.9*μ*_0_. **f**, Comparison of measured OD vs OD as expected from dilution. Actual measured OD (blue line) is shown compared to OD_*dilution*_(*t*) as estimated from eq. 28, once using the measured mean ribosome concentration (purple circles), and once using the fitted mean ribosome used in Supplementary Note 2.2 (dashed purple line). The comparison shows that the expected OD from a dilution-based process is 5-fold higher than the measured OD. **g**, Numerically calculated theoretical ribosome distributions. Four theoretical pdf-s (based on eq. 23), each having a different value of switching rate *γ_rib_*, but all having the same values for the rest of the model’s parameters (including *λ_a_*). For each of the density functions, a marker for its peak/peaks was plotted (black cross). The two density functions with the lower *γ_rib_* values showed bimodality while the two function with the higher *γ_rib_* values did not. **h**, A phase space picture showing a region of bimodality. A phase plot showing the value of the bimodality measure *d* (color-bar), calculated on theoretical densities (based on eq. 23) with varying values of switching rate *γ_rib_* and active degradation rate λ_*a*_. According to the parameters estimation (done in Supplementary Notes 2.1 and 2.2) the *S*.Tm population (*γ_rib_* ≈ 1.4*μ*_0_, λ_*a*_ ≈ 3.8*μ*_0_) lies in the region of binomdality (*d* > 0) while the *E. coli* population (*γ_rib_* ≈ 3*μ*_0_, λ_*a*_ ≈ 4.2*μ*_0_) lies in the non-bimodal region (*d* = 0). The two bacterial species are annotated with black circles and their uncertainty as error-bars.

**Extended Data Fig. 3:**
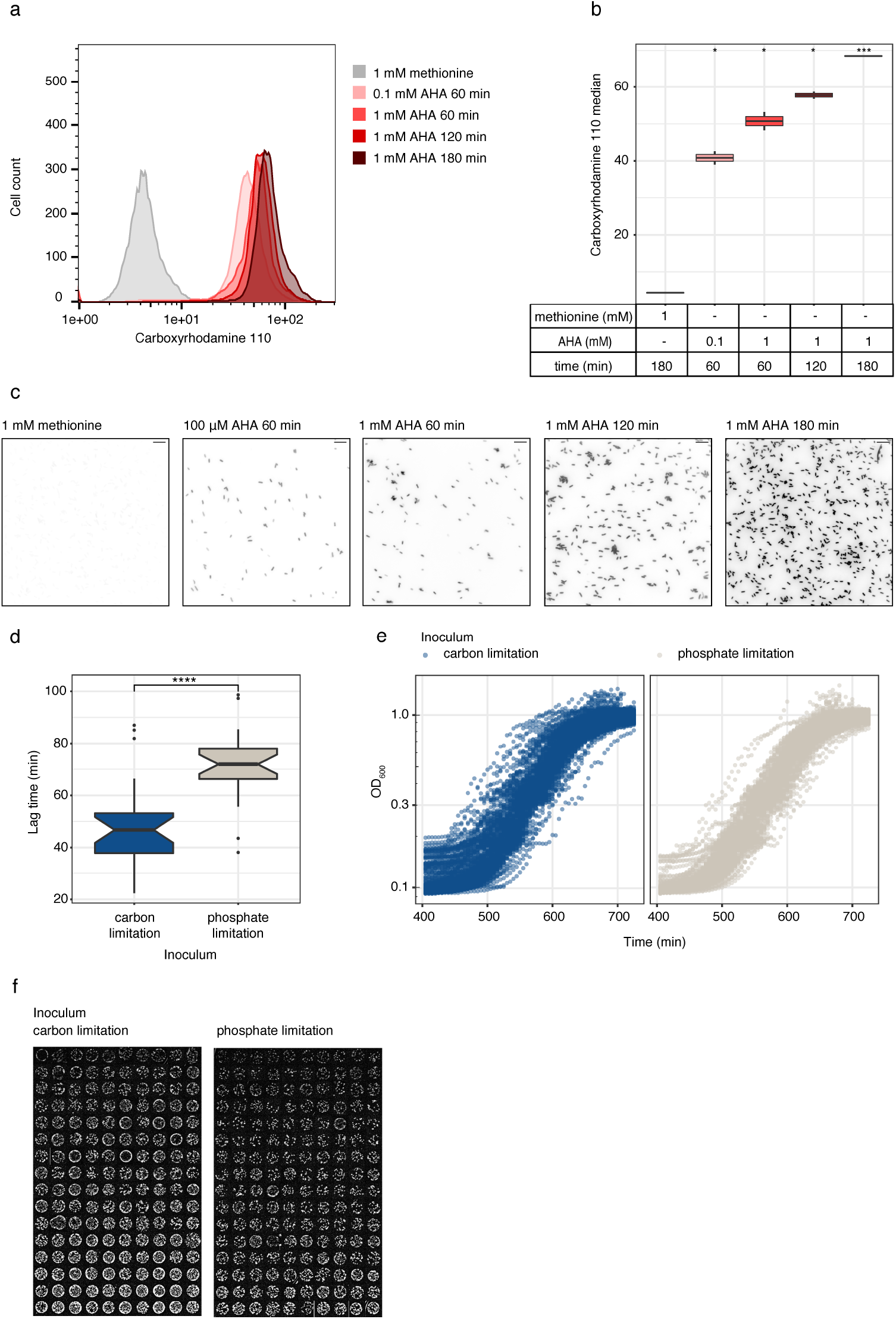
Quality controls of BONCAT. **a**, Density histograms of different BONCAT treatments showing AHA-carboxyrhodamine 110 incorporation levels (in shades of red) measured by flow cytometry. Methionine control is shown in grey (n = 10,000 cells). **b**, Boxplots of BONCAT treatments as in Fig. S3a, showing AHA-carboxyrhodamine 110 median. Unpaired t-test: *, *P* <0.05; ***, *P* <0.001. **c**, Representative microscopy images of BONCAT treatments as in Fig. S3a. Scale bar, 10 *μ*m. **d**, *S*.Tm cells inoculated from carbon-limited MM (in blue) display shorter lag time during nutritional upshifts compared to phosphate-limited (in grey). Nutritional upshifts curves shown in Fig. 3d were used to calculate lag times, shown as boxplots (n = 24). Unpaired t-test: ****, *P* <0.0001. **e**, Nutritional upshift curves obtained from single-sorted bacteria, used to calculate lag times in Fig.3e. Before the nutritional upshift, bacteria were grown in carbon- (left panel, in blue) or phosphate-limitation (right panel, in grey) (n = 96 – 114). **f**, Drop spotting following nutritional upshifts for carbon-limited (left panel, n = 160) and phosphate-limited (right panel, n = 160) cells. Images are sorted according to CFU counts.

**Extended Data Fig. 4:**
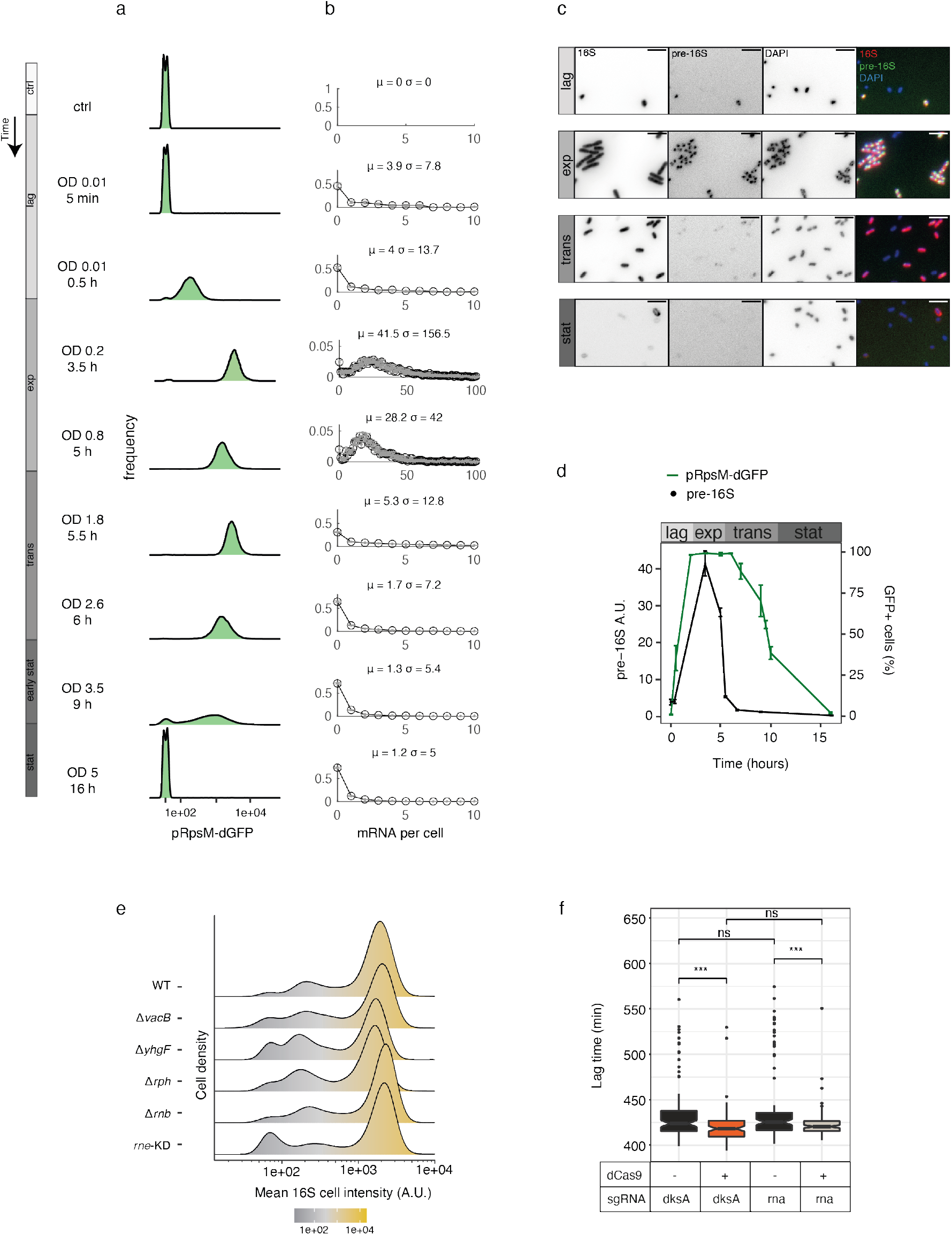
Comparison between pRpsM-dGFP reporter and pre-16S smFISH. **a**, Representative FACS histograms of *S*.Tm expressing dGFP under the control of the RpsM promoter from the indicated time points and OD_600_ (n = 10,000 cells). Non-fluorescent bacteria (ctrl) are shown as reference for background fluorescence. **b**, Distributions of total (per cell) pre-16S rRNA, from the indicated time points and OD_600_ during the growth curve. Means and standard deviations are shown. **c**, Representative microscopy images of 4 time points (lag, exponential, transition, and stationary) showing 16S rRNA (in red), pre-16S rRNA (in green), and DAPI (in blue). Single channels in black. Scale bar, 5 *μ*m. **d**, Transcription of rRNA and pRpsM-dGFP reporter expression during the different growth phases. rRNA transcription is presented as average of pre-16S rRNA molecules per cell (in black). pRpsM-dGFP activity is presented as % of dGFP+ cells (in green). **e**, Single cell nutritional upshifts from 4f, presented as boxplots. Unpaired t-test: ns, *P* >0.05; ***, *P* <0.001. **f**, Density histograms show 16S rRNA-FISH mean fluorescence intensity of individual *S*.Tm cells, from the indicated samples (n = 1538 – 1985 cells).

## Supplementary Notes

### 1 Model for ribosome dynamics

Our main goal here is to theoretically address the observed ribosome concentration dynamics described in the main text. We focus on two main qualitative features: the gradual decrease of ribosome concentration over time and the emergence of bimodal concentrations distribution across the cell population.

We will analyze two models. The first assumes a dilution-based process at which the transcription of new ribosomes is terminated. Ribosome concentration is halved with every population doubling (as the total number of ribosomes is fixed while the total number of cells increases). The second model expands the first by adding an *active-degradation* mechanism, in which certain proteins arise in the cell, responsible for degrading ribosomes. As we show in Supplementary Note 3, the dilution-based model makes predictions order of magnitude far from measured results and plays a role here as a logical step towards the more complex model that includes an *active-degradation* mechanism.

#### 1.1 Exponential growth dynamics

In order to approach the observed dynamics analytically, will adapt the gene expression model of [1]. We first consider the number of proteins in all cells (*P_i_*) at a given time. Its rate of change is then given by:

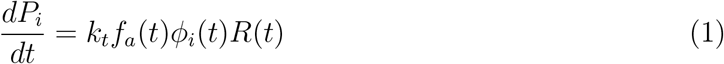

where *R*(*t*) is the mean ribosome number in a cell, *k_t_* is the translation rate per ribosome of a typical mRNA, assumed to be constant, *ϕ_i_*(*t*) is the gene allocation fraction of protein *i* (*i.e.* the fraction of RNAPs that are working on protein *i*’s gene), and *f_a_* is the fraction of ribosomes active for translation.

For the special case of *R*(*t*), eq. 1 reads:

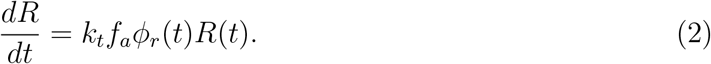

We estimate the cell volume *V*(*t*) as proportional to the sum of all proteins, namely - *V*(*t*) = *α* ∑ *P_i_*(*t*), having *α* as the mean volume a single protein occupies [1–4]. This leads to:

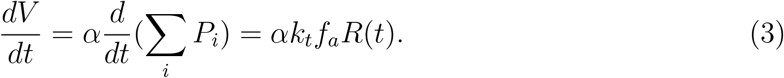

Now we can write the deterministic dynamics for the ribosome concentration 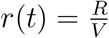

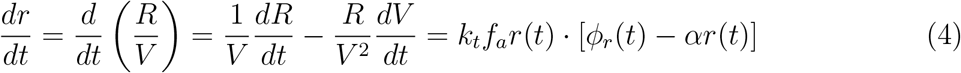

with a stable fixed point at 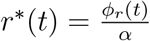.

In an assumed steady-state exponential growth, we expect the allocation fraction *ϕ_r_* to be the steady-state ratio of the R-protein mass out of the total protein mass [5]. Hence, we approximate *ϕ_r_* as constant during exponential growth, namely - 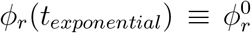, suggesting a constant mean ribosome concentration - 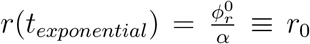. Following the same reasoning, in exponential growth, we expect a constant value of growth rate *μ*_0_ supporting the exponential trend of cell number *N*(*t*) ∝ *e*^*μ*_0_*t*^. It is also assumed that the fraction of active ribosomes is constant in exponential growth, namely - 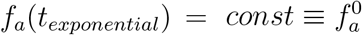. Finally we define 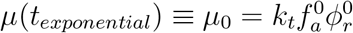.

#### 1.2 Stochasticity in switching times

An essential aspect of both models is the assumption that at some point in time *t*_0_, cells detect that nutrients in the environment are bordering depletion and gradually switch from a state of exponential growth to a state where new ribosome transcription is terminated. This time refers to the “transition phase” found in the main text. Moreover, in our current setup, we also suggest that the process of switching is a stochastic one, in which a cell has a probability of switching in a time gap δ*t* of *γ_rib_δt*, where *γ_rib_* is a constant switching rate. This makes Δ*t* - the time between *t*_0_ and the time at which a cell switches 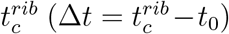, an exponentially distributed random variable. Its probability density function (pdf) is given by:

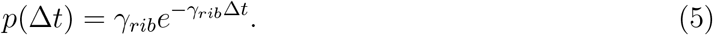

In the following sections, we will be interested in the expression of probability densities of the ribosome concentration r(t). For that, we will derive a relation between Δ*t*, *r*(*t*) and *t*, and use a change of variables to get an expression for *p*(*r, t*).

#### 1.3 Dilution based process

In this model, we look at a cell, assumed to start at *t* = 0 in a steady state exponential growth, with *r*(0) = *r*_0_. The cell is assumed to switch off its ribosome transcription at 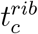. We model it by letting the ribosome genetic allocation fraction *ϕ_r_*(*t*) drop to zero at the critical time. We thus have:

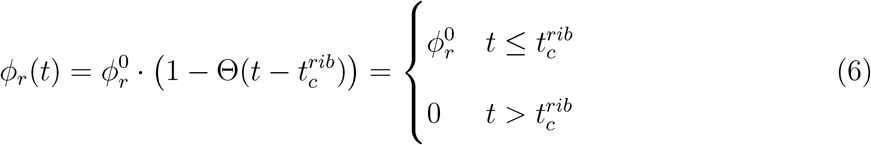

where Θ is the Heaviside step function. In the current modeling of a dilution process, we also assume that the ribosomes are well mixed in the cell. Thus with each division of a mother cell, the two daughter cells receive an equal number of ribosomes (both active and inactive).

Plugging the above into eq. 2 leads to:

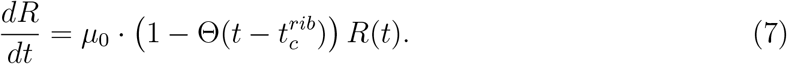

Solving, we find:

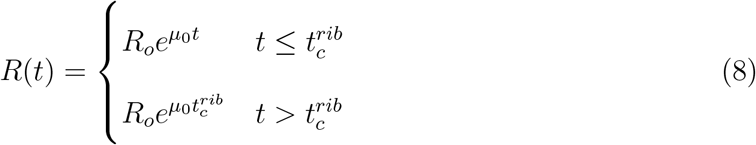

where *R*_0_ is an arbitrary initial condition that follows the steady state assumption, namely that 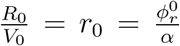, with *V*_0_ being the initial condition for the volume. We do the same procedure for the volume equation (3) to get:

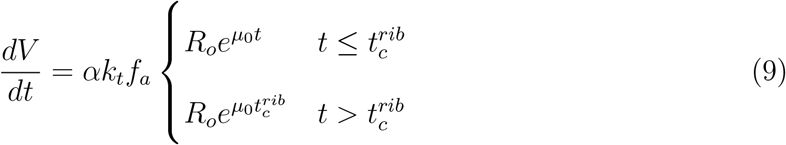

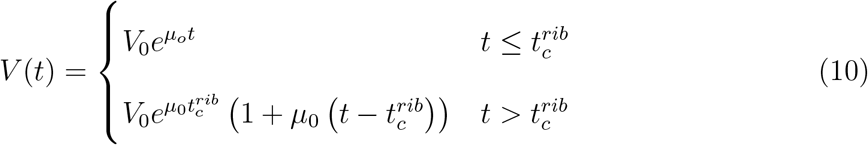

We finally put everything together to find the expression for the ribosome concentration:

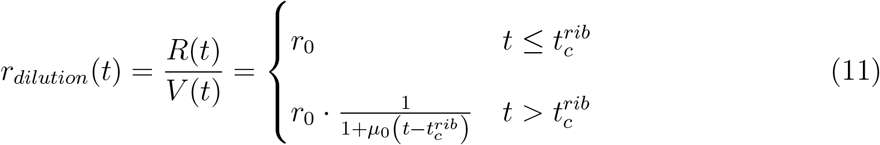

We then look at a collection of cells, each obeying the above dynamics but differs by its switching time - 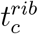, which is now considered a random variable that is distributed exponentially. Consequently, *r*(*t*) is a random variable for which we want to find the probability density function. We assume that the entire cell population starts from the steady state concentration *r*_0_, and so for times *t* < *t*_0_, we have a peaked distribution *p*(*r, t*) = *δ*(*r* – *r*_0_), where *δ*() is the Dirac delta function. In later times *t* > *t*_0_, we invert the relation shown in eq. 11 to get an expression for 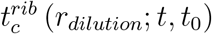 and Δ*t* (*r_dilution_*; *t*, *t*_0_) (with 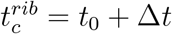). We then use a change of variables to get an expression for *p*(*r, t*):

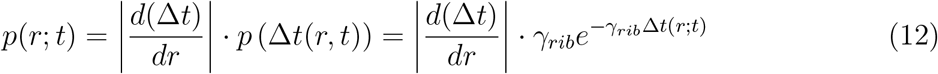

where we substituted *p*(Δ*t*) with equation 5. Specifically using the resulting Δ*t_dilution_*(*r, t*)

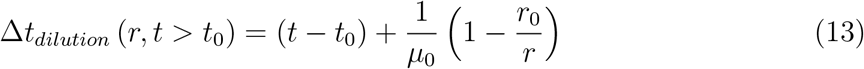

in eq. 12, the pdf reads:

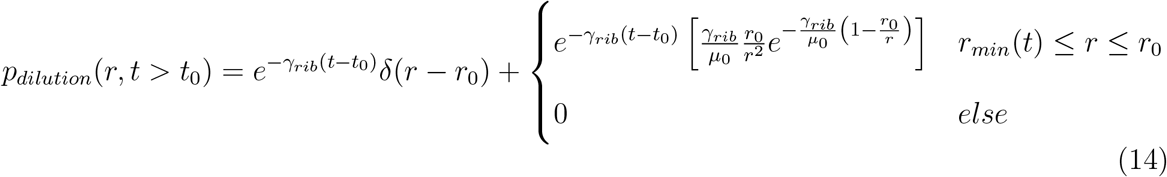

where *r_min_*(*t*) is the minimal ribosome concentration possible given by the case where the switching time is exactly *t*_0_, namely - 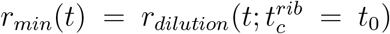. This results in a probability density function comprised of two distributions, one (the first term) being the portion of the population which hasn’t switched and has a fixed ribosome concentration, while the other (second term) being the portion of switched cells, whose ribosome concentration is gradually decreasing.

#### 1.4 Active degradation based process

As suggested in the main text and we will show in Supplementary Note 3 the dilution process alone is insufficient to explain the data. We next turn to consider a case where in addition to the transcription termination assumed before, an active degradation mechanism starts degrading ribosomes at a constant rate λ_*a*_. We further assume that this new mechanism degrades mainly active ribosomes rather than inactive ribosomes [6]. The presence of ribosomes that do not get degraded facilitates a stable lower ribosomes concentration state, which is also apparent in the experimental data at longer times (whose value is ~ 2% of the measured r0).

For our model, in addition to the assumptions taken in Supplementary Note 1.3, we also assume that the degradation mechanism degrades active ribosomes only, with a rate of λ_*a*_ (*R*(*t*) – *R_in_*(*t*)), while the inactive ribosomes (*R_in_*(*t*) = (1 – *f_a_*(*t*))*R*(*t*)) are not affected. This means that once a cell has switched, the number of inactive ribosomes remains constant, namely -

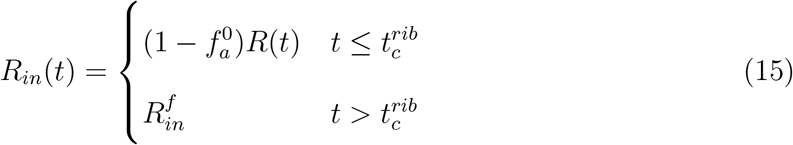

where we defined 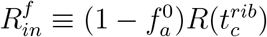.

We can now write the rate equations for both the ribosome number and the volume:

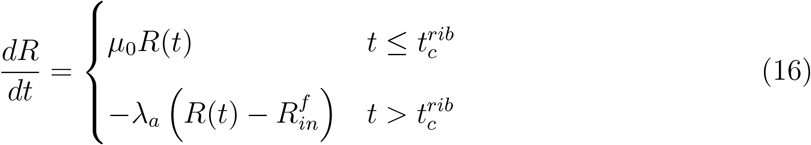

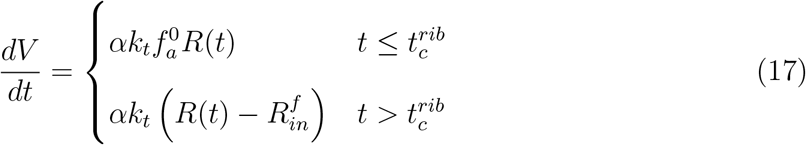

where we assumed that the degraded ribosomes do not decrease the cell volume, as their components remain part of the cell dry mass [4]. Following the same steps as done previously (Supplementary Note 1.3), one can get an expression for the ribosome concentration:

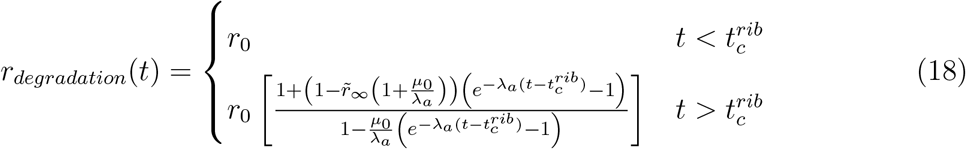

where we defined 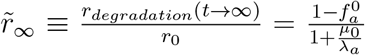, which represents the low steady-state ribosome concentration in long times (relative to *r*_0_). As before, we will invert the above relation to get Δ*t_degradation_*(*r,t*) and plug it into eq. 12 to obtain:

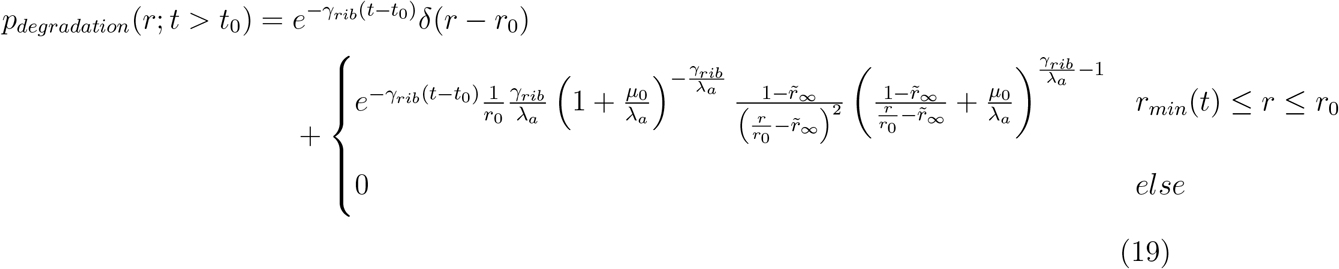

where 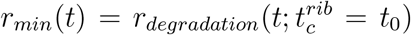. As for the dilution-only case, when adding active degradation, we get a composition of a peak at *r*_0_ representing the exponentially growing cells and a second broader peak representing the switched cells for which the ribosome concentration is decaying. It is also useful to define the mean ribosome concentration as follows:

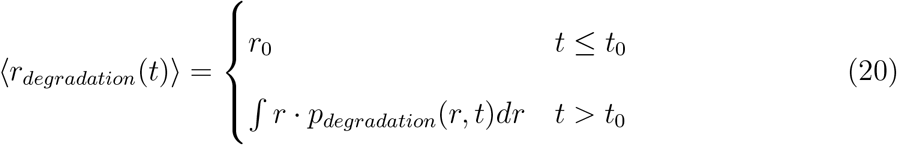

#### 1.5 Multiplicative noise

In order to analyze the experimental results, we assume that during exponential growth, cells are in a steady state with a fixed ribosome concentration (*r*(*t_exponential_*) = *r*_0_). Thus, we expect to see a peak in the ribosome distribution at *r*_0_ (i.e., *p*(*r, t_exponential_*) = *δ*(*r* – *r*_0_)). In reality, there is usually phenotypic variability within a population [7–9]. However, the current experimental results showed that the variability of *r*_0_ (*CV* ~ 30%) was much larger than expected from phenotypic variability (*CV* ~ 5%)[10], indicating that another source of noise is dominating. Examining the lower ribosome peak, we noted that its CV was approximately 50%. This contradicts the notion that the observed noise is additive, as an additive noise would result in a CV of approximately 400% - eight times higher than the observed value. We therefore characterized the noise as a Speckle multiplicative noise, as was previously reported to be present in fluorescence microscopy experiments [11].

The noise was formulated by defining 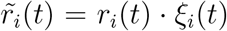 for a cell *i* at time *t*, Where *ξ_i_* is an independent random variable that represents Speckle noise, commonly modeled using the Gamma distribution [12] with pdf

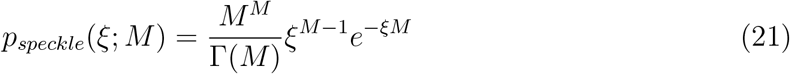

where 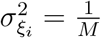, and 〈*ξ_i_*〉 = 1. Given 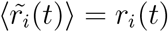, the signal-to-noise ratio (SNR) follows:

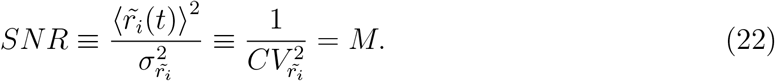

Later in our analysis, it will be important to assess the noise level in the measured *r*_0_ value and calculate its signal-to-noise ratio (SNR), thus finding *M* (again naively assuming that *p*(*r, t_exponential_*) = *δ*(*r* – *r*_0_)). Finally, considering the the distribution of 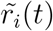 across cells population, we get the following expression:

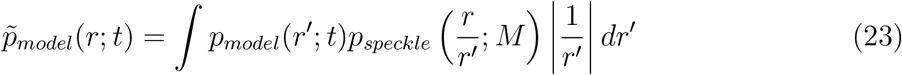

where *p_model_*(*r, t*) represents either the pdf for the dilution-only case (eq. 14) or the pdf that considers active degradation (eq. 19)

#### 1.6 Bimodality and alternative models

he formation of a bimodal distribution is often attributed to the presence of two distinct subpopulations with different characteristics [13]. The current experimental results suggest the presence of such two subpopulations, one with high ribosome levels and the other with low ribosome levels, resulting in a ribosome pdf that is a mixture of two distributions, each representing its respective subpopulation. At the initial stages of the experiment, all cells are growing exponentially, and the population is homogeneous, characterized by the steady-state ribosome level *r*_0_. Hence, at some point, cells “decide” if and when they should switch to the subpopulation with lower ribosome levels. In the current work, this “decision” made by portion of the population is considered a stochastic process and can be modeled in various ways.

Within our current theoretical framework, we modeled the process using stochastic switching, in which cells transition from an exponential growth state to a state of ribosome transcription shut-down, characterized by the onset of constant ribosome degradation. This transition occurs gradually over time, with a randomly determined switching time. As result of this process, a subpopulation of cells is formed with high levels of ribosomes and another subpopulation with declining levels of ribosomes.

Nevertheless, other alternative models exist, capable of supporting the observed bi-modality. One possibility, for instance, is that all cells simultaneously transition to a non-transcribing state at time *t*_0_. In this scenario, each cell would have a randomly assigned degradation rate λ_*a*_ drawn from a distribution. A bimodal distribution of degradation rates, which could be a result of a bistable circuit [14], would, in turn, lead to a bimodal distribution of ribosomes, with one subpopulation rapidly degrading and the other degrading slowly. The bimodality in this case is manually induced rather than arising from the underlying dynamics as it does in the previous model (which we will refer to as the *degradation model*). Regardless, based on the available experimental results, we cannot eliminate this scenario at present. However, it can be investigated further in future studies by monitoring ribosomes in individual cells over time to determine the distribution of degradation rates.

Another noteworthy variant, which aligns more closely with the *degradation model*, involves the onset of ribosome degradation occurring at a later time after transcription has ceased. Specifically, the model proposes that cells first shut down transcription (for simplicity, all simultaneously), then each cell has a random delay before the onset of ribosome degradation. This attempts to simulate a rapid detection of nutrient depletion by the population and a slower cascade of biological events that triggers the activation of ribosome degradation.

Finally, there is a characteristic in the data that is not accounted for by the *degradation model*. This feature is a gradual decrease in the ribosome concentration of the high-ribosome subpopulation. While the two proposed alternatives provide a qualitative explanation for this decrease, a closer examination reveals that the predicted decay rate should be equal to or faster than a dilution decay rate. By comparing the observed decay with the theoretical dilution decay, it becomes evident that the decrease seen in the data is slower than expected from dilution, thus rendering the alternatives less compelling.

### 2 Estimation of model parameters

In this section, we use experimental data to estimate the parameters of our theoretical model. We test our model against two experimental data sets, one from *S*.Tm and the other from *E. coli*. The results of this parameter estimation process will provide insights, in following sections, into how well the current model describes the underlying mechanisms of phenotypic switching and degradation during starvation and the transition to stationary growth phase.

As mentioned, every analysis that will follow in this section is based on experiments that used *S*.Tm and *E. coli*. Both experiments sampled the cells’ fluorescence intensity from different probes, as elaborated on in the main text. The two aforementioned experiments contain 13 samples at different time points. The first 4 time points were taken when the population was presumably in an exponential growth phase, while the other 9 are in the transition to stationary and stationary phase. Additionally, for each time point of the samples mentioned above, a measure of the population’s OD was taken.

For our analysis, we used the fluorescence data from the rRNA 16S probe. It is assumed throughout the analysis that the rRNA 16S fluorescence concentration is proportional to the ribosome concentration. As we are primarily interested in relative ribosome concentration, we can ignore proportionality and use ribosome concentration and fluorescence concentration interchangeably. For each sample, the background intensity was subtracted and cells with intensities exceeding the 99.5% percentile were excluded as outliers.

For estimating the growth rate *μ*_0_, a fit was made using the simple exponential growth equation -

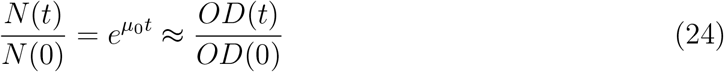

where we assumed the number of cell *N* is proportional to the *OD* measure, *N*(*t*) ∝ *OD*(*t*). The fitting was done on the first 4 OD samples taken when the population grew exponentially. We found that for *S*.Tm 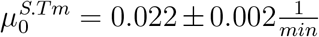 and for *E. coli* 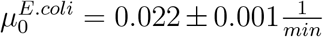.

#### 2.1 Estimating the switching rate *γ_rib_*

To estimate the switching rate *γ_rib_* independently of the active degradation rate λ_*a*_, we turn to observe the dynamics of the number of exponentially growing cells *N_exp_*(*t*), before and after the nutrient depletion tipping point at *t*_0_. We again use *OD*(*t*) as a proxy for the number of cells, thus having *N_exp_*(*t*) = *f_exp_*(*t*)*N*(*t*) ∝ *f_exp_*(*t*)*OD*(*t*) ≡ *OD_exp_*(*t*), where *f_exp_*(*t*) is the fraction of exponentially growing cells at time *t*. This leads to:

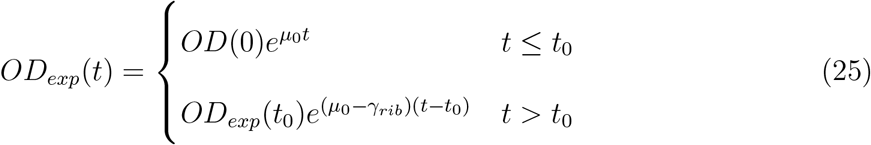

Next, we estimate *OD_exp_*(*t*) from the data. To do so, we will estimate the values of *f_exp_*(*t*) and multiply by the measured *OD*(*t*). For each time sample, we consider the higher ribosome peak as representing the exponential cells. The peak is fitted by a curve (according to a Gamma distributed noise as discussed in Supplementary Note 1.5). The area of the fitted peak is then considered *f_exp_*(*t*), and its error is estimated by the fitting error. Finally, using the estimated *OD_exp_*(*t*) and eq. 25, we fit both the switching rate *γ_rib_* and the tipping time *t*_0_ (Fig. N. 1).

**Fig. N. 1:**
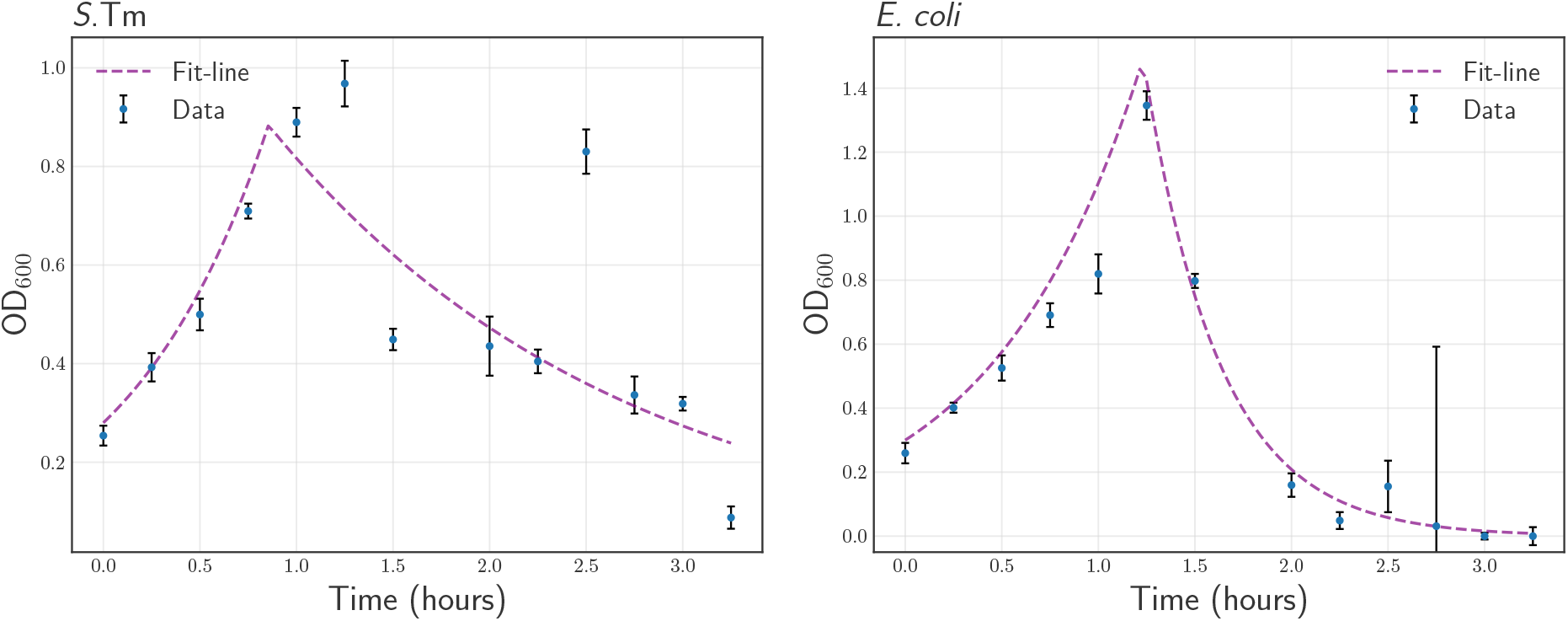
*OD_exp_*(*t*) fitting using eq. 25. A fit of the theoretical expression eq. 25 (dashed purple line) to the measured *OD_exp_*(*t*) (blue circles) for the two bacteria types. For *S*.Tm, the fitting parameters came as *γ_rib_* = 1.4*μ*_0_ ± 0.1*μ*_0_ and *t*_0_ = 0.85 ± 0.08*hr*. For *E. coli*, the fitted parameters are *γ_rib_* = 3.0*μ*_0_ ± 0.2*μ*_0_ and *t*_0_ = 1.23 ± 0.02*hr*.

#### 2.2 Estimating the active degradation rate *λ_a_*

We aim to compare our model’s pdf (eq. 19) with experimental ribosome concentration data. However, a direct comparison is challenging due to high measurement noise and model limitations. Despite these limitations, our model may still provide reasonable estimates of the distribution mean at a given time. We can compare the theoretical mean 〈*r_degradation_*(*t*)〉 (eq. 20) with the average ribosome concentration over the population at various time points and fit for the active degradation rate λ_*a*_.

In order to proceed, we must determine two parameters required by the model. The first parameter, *r*_0_, is calculated as the average of the centers of the high-ribosome peaks (as described in Supplementary Note 2.1) across the various time points. The second parameter, 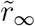, represents the low steady-state ribosome concentration in long times, and is estimated in a similar manner as *r*_0_ by analyzing only the lower ribosome peak.

Finally, we use the above to fit eq. 20 to the data, as previously discussed (Extended Data Fig. 2c). We were able to obtain a reasonable fit, which estimates the amount of active degradation needed to reduce the average ribosome value by an order of magnitude during the course of the experiment.

### 3 Comparing dilution and active-degradation based models

A main concern in the current work is whether an active degradation mechanism is involved in the process of ribosome concentration decay. The suggested alternative was a dilution-based process. The first analytical evidence for active degradation presence was presented in Supplementary Note 2.2 when fitting the mean ribosome concentration over time suggested that the strictly non-zero active degradation rate (λ_*a*_ ≈ 4*μ*_0_) is needed to get a good fit. In this section, we provide an alternative approach that will further strengthen the above conclusion and will argue against a dilution-only process. In a dilution-based process, the mean ribosome concentration 〈*r*(*t*)〉 is approximately halved in each division of the population, so one may approximate-

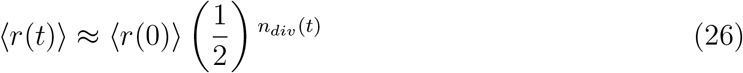

where *n_div_*(*t*) is an approximated number of division taken in time *t*. Given the number of population divisions and the initial cell number, one can get the number of cells at a time point, namely -

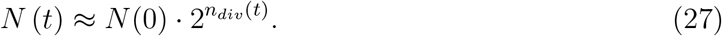

Now, using the assumed proportionality of the number of cells and the OD, one can plug the expression for *n_div_*(*t*) (from eq. 26) and plug it into eq. 27 to get

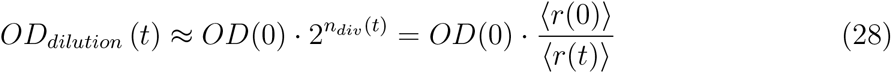

We now turn to compare the above with the actual OD measure. We plot the expected result from eq. 28 once using the measured mean ribosome concentration and once using the analytical expression (eq. 20) as was fitted in section 2.2 (Extended Data Fig. 2d). Observing the resulting comparison, it is clear that there is a significant gap (~ 5 fold) between the measured OD and the one expected from dilution (eq. 28). This gap supports our conclusion that a dilution-only processes is inadequate.

### 4 The conditions for bimodality to emerge

An interesting aspect of observed ribosome dynamics is the emergence of bimodal ribosome distributions. As discussed in the main text, bimodality could facilitate a form of adaptation in specific environments. Furthermore, as mentioned in the main text, not all tested bacteria species showed bimodal trends. In particular, *S*.Tm showed a bimodal trend (Fig. 1d), while *E. coli* did not (Fig 2b).

Within the framework of our current model concerning ribosome transcription switching and active degradation of ribosomes, we claim that the emergence of bimodality is tightly dependent on the two main model’s parameters - the switching rate *γ_rib_* and the active degradation rate *λ_a_* (both measured relative to the growth rate *μ*_0_).

To gain a deeper understanding of this parametric dependence, we calculate the density function 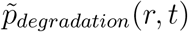 according to eq. 23, with varying values of *γ_rib_* and *λ_a_*, and examine the presence of bimodality for each value combination. It is worth noting that the level of experimental noise also influences the presence of observable bimodality since higher noise levels make two distribution peaks harder to distinguish. Next, we examine the probability density functions at the time in which half of the cells have switched - 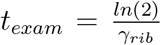, and the distribution is expected to be roughly in its maximal variance. We determine and fix the additional parameters needed by the pdf 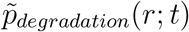, including the noise coefficient of variation, 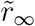, *r*_0_, and *μ*_0_. The noise coefficient of variation represents the experimental fluorescence noise (as was described in Supplementary Note 1.5) and was estimated from the data (*CV* ≈ 30%). In addition 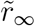 was also set in accordance with data estimations 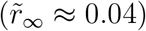, while *r*_0_ and *μ*_0_ were set to 1. This setting of *r*_0_ = *μ*_0_ = 1 should not alter the shape, and thus the bimodality of the distribution, as the expression 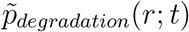 only contains rates relative to *μ*_0_ and concentration values (*r*) relative to *r*_0_.

As an example, we plotted four 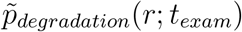 distributions with varying *γ_rib_* values (Extended Data Fig. 2e) and a fixed *λ_α_*. In addition to the distributions themselves, we marked each of the peak/peaks in each distribution. The two distributions with the higher *γ_rib_* values have only one observable peak (i.e., not bimodal). In comparison, the two distributions with the lower valued *γ_rib_* possess two observable peaks (i.e., bimodal).

Next, we quantify bimodality by defining the distance between peaks *d* as the measure of bimodality. This distance d is defined as zero when only one peak is present (i.e., no bimodality). Making use of this measure, we run a numerical analysis that calculates d for different 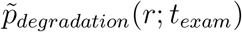 distributions, each corresponding to varying values of *γ_rib_* and *λ_a_*. Finally we plot the *γ_rib_* - *λ_a_* phase space picture and the measure of bimodality (d) for each (*γ_rib_, λ_a_*) combination (Extended Data Fig. 2f). The phase space contains a region of bimodality (*d* > 0) and a region of non-bimodality (*d* = 0). It is interesting to note that according to the parameters we estimated in Supplementary Notes 2.1 and 2.2, *S*.Tm lies deep within the bimodal region (*γ_rib_* ≈ 1.4*μ*_0_, λ_*a*_ ≈ 3.8*μ*_0_) while *E. coli* lies mostly in the non-bimodal region (*γ_rib_* ≈ 3*μ*_0_,λ_*a*_ ≈ 4.2*μ*_0_). This is consistent with the aforementioned experimental observations, in which *S*.Tm population display bimodality while *E. coli* does not (Figs. 1d and 2b).

It should be noted that our model will always generate bimodal distributions in the absence of any experimental noise (as depicted in Fig. N. 2).

**Fig. N. 2:**
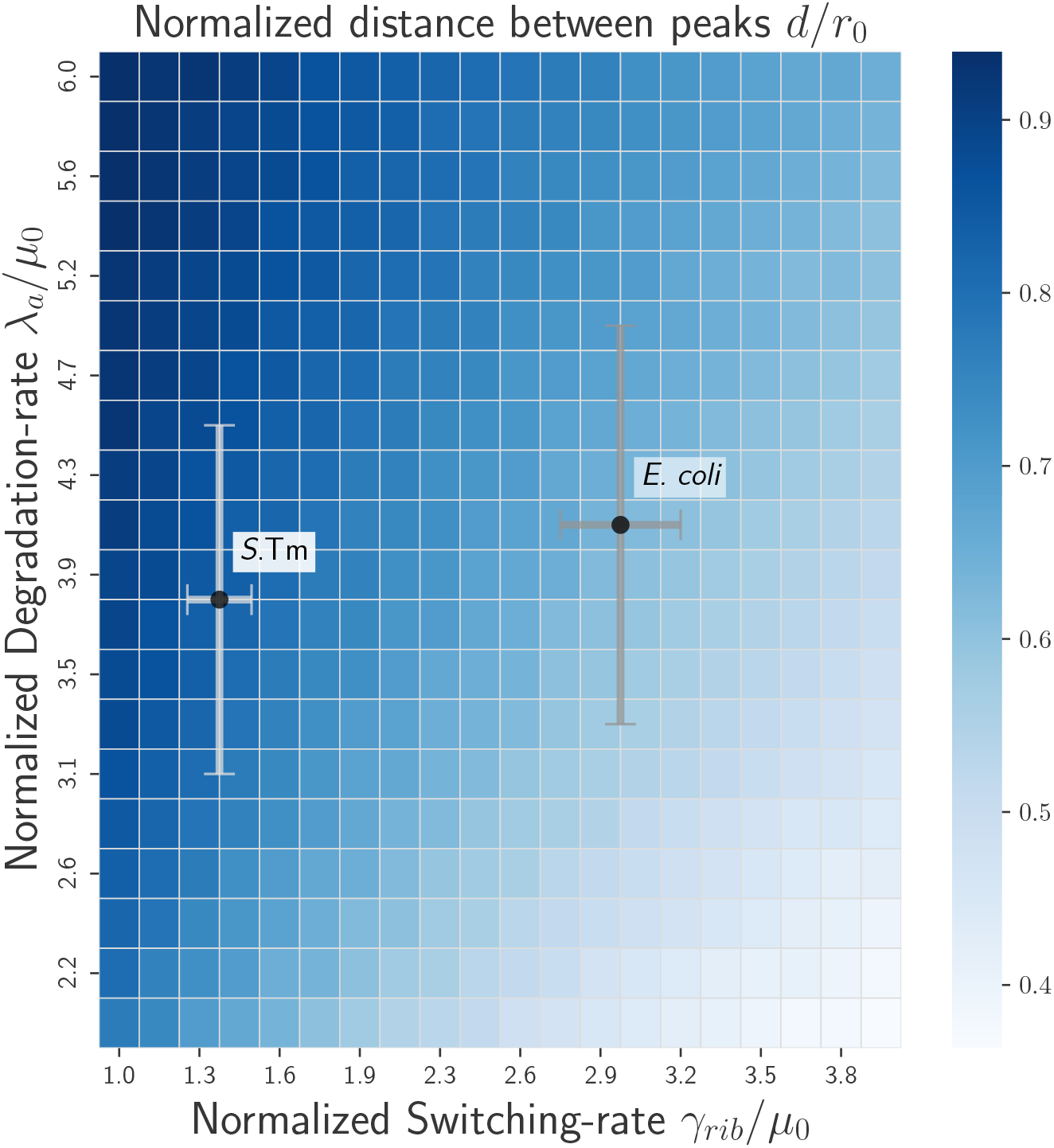
A phase space picture without experimental noise. A phase plot showing the value of the bimodality measure *d* relative to *r*_0_ (color-bar), similar to Extended Data Fig. 2f only calculated using theoretical densities clean of experimental noise (eq. 19). The *d/r*_0_ values are 0.83 for *S*.Tm and 0.62 for *E. coli*.

Hence, the discussion about the presence of bimodality in various bacteria pertains to the extent to which bimodality can be observed under a specific level of noise. Our findings suggest that, even in the absence of experimental noise, *S*.Tm might demonstrate a higher degree of bimodality than *E. coli*, as the calculated bimodality measure d was approximately 27% higher for *S*.Tm.

Finally, the phase space in Extended Data Fig. 2f provides us with information about the conditions for bimodality in a dilution-only process (with no active degradation, λ_*a*_ = 0). By analyzing the critical parameter values 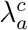 and 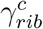 (which separate the bimodal and non-bimodal regions), we use a linear fit 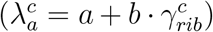 to extrapolate the maximal switching rate supporting bimodality in the absence of degradation, namely 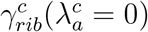. This maximal switching rate was estimated as 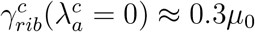, which is an order of magnitude smaller than the estimated switching rates (~ 1.4*μ*_0_ – 3*μ*_0_), suggesting that dilution alone cannot explain the observed bimodality in our current theoretical framework.

## References

1. Röder, J., Felgner, P. & Hensel, M. Comprehensive Single Cell Analyses of the Nutritional Environment of Intracellular Salmonella enterica. Frontiers in Cellular and Infection Microbiology 11, 83 (2021).

2. Deutscher, M. P. Degradation of Stable RNA in Bacteria. Journal of Biological Chemistry 278, 45041–45044 (2003).

3. Piir, K., Paier, A., Liiv, A., Tenson, T. & Maiväli, Ü. Ribosome degradation in growing bacteria. EMBO reports 12, 458–462 (2011).

4. Scott, M., Gunderson, C. W., Mateescu, E. M., Zhang, Z. & Hwa, T. Interdependence of Cell Growth and Gene Expression: Origins and Consequences. Science 330, 1099–1102 (2010).

5. Zundel, M. A., Basturea, G. N. & Deutscher, M. P. Initiation of ribosome degradation during starvation in Escherichia coli. RNA (2009).

6. Gausing, K. Regulation of ribosome production in Escherichia coli: Synthesis and stability of ribosomal RNA and of ribosomal protein messenger RNA at different growth rates. Journal of Molecular Biology 115, 335–354 (1977).

7. Mandelstam, J. THE INTRACELLULAR TURNOVER OF PROTEIN AND NUCLEIC ACIDS AND ITS ROLE IN BIOCHEMICAL DIFFERENTIATION. Bacteriological Reviews 24, 289–308 (1960).

8. Maruyama, H. & Mizuno, D. Ribosome degradation and the degradation products in starved Escherichia coli. I. Comparison of the degradation rate and of the nucleotide pool between Escherichia coli B and Q-13 strains in phosphate deficiency. BBA Section Nucleic Acids And Protein Synthesis (1970).

9. Ben-Hamida, F. & Schlessinger, D. Synthesis and breakdown of ribonucleic acid in Escherichia coli starving for nitrogen. BBA Section Nucleic Acids And Protein Synthesis (1966).

10. Jacobson, A. & Gillespie, D. Metabolic events occurring during recovery from prolonged glucose starvation in Escherichia coli. Journal of bacteriology (1968).

11. McCarthy, B. J. The effects of magnesium starvation on the ribosome content of Escherichia coli. BBA Specialized Section on Nucleic Acids and Related Subjects (1962).

12. Kaplan, R. & Apirion, D. The fate of ribosomes in Escherichia coli cells starved for a carbon source. Journal of Biological Chemistry (1975).

13. Basturea, G. N., Zundel, M. A. & Deutscher, M. P. Degradation of ribosomal RNA during starvation: comparison to quality control during steady-state growth and a role for RNase PH. RNA (New York, N.Y.) 17, 338–345 (2011).

14. Sulthana, S., Basturea, G. N. & Deutscher, M. P. Elucidation of pathways of ribosomal RNA degradation: an essential role for RNase E. RNA 22, 1163 (2016).

15. Prossliner, T., Gerdes, K., Sørensen, M. A. & Winther, K. S. Hibernation factors directly block ribonucleases from entering the ribosome in response to starvation. Nucleic Acids Research 49, 2226–2239 (2021).

16. Villadsen, I. S. & Michelsen, O. Regulation of PRPP and nucleoside tri and tetraphosphate pools in Escherichia coli under conditions of nitrogen starvation. Journal of bacteriology 130, 136–143 (1977).

17. Metzger, S., Schreiber, G., Aizenman, E., Cashel, M. & Glaser, G. Characterization of the relA1 mutation and a comparison of relA1 with new relA null alleles in Escherichia coli. Journal of Biological Chemistry 264, 21146–21152 (1989).

18. Gentry, D. R. & Cashel, M. Mutational analysis of the Escherichia coli spoT gene identifies distinct but overlapping regions involved in ppGpp synthesis and degradation. Molecular microbiology 19, 1373–1384 (1996).

19. Ryals, J., Little, R. & Bremer, H. Control of rRNA and tRNA syntheses in Escherichia coli by guanosine tetraphosphate. Journal of Bacteriology 151, 1261 (1982).

20. Spira, B., Silberstein, N. & Yagil, E. Guanosine 3’,5’-bispyrophosphate (ppGpp) synthesis in cells of Escherichia coli starved for Pi. Journal of Bacteriology 177, 4053–4058 (1995).

21. Iyer, S., Le, D., Park, B. R. & Kim, M. Distinct mechanisms coordinate transcription and translation under carbon and nitrogen starvation in Escherichia coli. Nature Microbiology 3, 741–748 (2018).

22. Potrykus, K. & Cashel, M. (p)ppGpp: still magical? Annual review of microbiology 62, 35–51 (2008).

23. Sanchez-Vazquez, P., Dewey, C. N., Kitten, N., Ross, W. & Gourse, R. L. Genome-wide effects on Escherichia coli transcription from ppGpp binding to its two sites on RNA polymerase. Proceedings of the National Academy of Sciences of the United States of America 116, 8310–8319 (2019).

24. Paul, B. J. et al. DksA: A critical component of the transcription initiation machinery that potentiates the regulation of rRNA promoters by ppGpp and the initiating NTP. Cell 118, 311–322 (2004).

25. Aiso, T., Yoshida, H., Wada, A. & Ohki, R. Modulation of mRNA stability participates in stationary-phase-specific expression of ribosome modulation factor. Journal of bacteriology 187, 1951–1958 (2005).

26. Yamagishi, M. et al. Regulation of the Escherichia coli rmf gene encoding the ribosome modulation factor: growth phase-and growth rate-dependent control. The EMBO journal 12, 625–630 (1993).

27. Izutsu, K., Wada, A. & Wada, C. Expression of ribosome modulation factor (RMF) in Escherichia coli requires ppGpp. Genes to cells: devoted to molecular cellular mechanisms 6, 665–676 (2001).

28. Shimada, T. et al. Classification and strength measurement of stationary-phase promoters by use of a newly developed promoter cloning vector. Journal of bacteriology 186, 7112–7122 (2004).

29. Durfee, T., Hansen, A. M., Zhi, H., Blattner, F. R. & Ding, J. J. Transcription profiling of the stringent response in Escherichia coli. Journal of bacteriology 190, 1084–1096 (2008).

30. Prossliner, T., Skovbo Winther, K., Sørensen, M. A. & Gerdes, K. Ribosome hibernation 2018.

31. Arrigucci, R. et al. FISH-Flow, a protocol for the concurrent detection of mRNA and protein in single cells using fluorescence in situ hybridization and flow cytometry. Nature Protocols 2017 12:6 12, 1245–1260 (2017).

32. Skinner, S. O., Sepúlveda, L. A., Xu, H. & Golding, I. Measuring mRNA copy number in individual Escherichia coli cells using single-molecule fluorescent in situ hybridization. Nature Protocols (2013).

33. Heo, M. et al. Impact of fluorescent protein fusions on the bacterial flagellar motor. Scientific Reports 2017 7:1 7, 1–10 (2017).

34. Yang, S., Veide, A. & So, E. Proteolysis of fusion proteins: stabilization and destabilization of staphylococcal protein A and Escherichia coli beta-galactosidase. Biotechnology and Applied Biochemistry 22, 145–159 (1995).

35. Gray, W. T. et al. Nucleoid Size Scaling and Intracellular Organization of Translation across Bacteria. Cell (2019).

36. Weng, X. et al. Spatial organization of RNA polymerase and its relationship with transcription in Escherichia coli. Proceedings of the National Academy of Sciences of the United States of America (2019).

37. Hsu, D., Shih, L. M. & Yuan Chung Zee. Degradation of rRNA in Salmonella strains: a novel mechanism to regulate the concentrations of rRNA and ribosomes. Journal of Bacteriology 176, 4761–4765 (1994).

38. Li, S. H. J. et al. Escherichia coli translation strategies differ across carbon, nitrogen and phosphorus limitation conditions. Nature Microbiology 2018 3:8 3, 939–947 (2018).

39. Dieterich, D. C., Link, A. J., Graumann, J., Tirrell, D. A. & Schuman, E. M. Selective identification of newly synthesized proteins in mammalian cells using bioorthogonal noncanonical amino acid tagging (BONCAT). Proceedings of the National Academy of Sciences of the United States of America 103, 9482–9487 (2006).

40. Cain, A. K. et al. A decade of advances in transposon-insertion sequencing. Nature Reviews Genetics 2020 21:9 21, 526–540 (2020).

41. Bernadac, A., Gavioli, M., Lazzaroni, J. C., Raina, S. & Lloubès, R. Escherichia coli tol-pal Mutants Form Outer Membrane Vesicles. Journal of Bacteriology 180, 4872 (1998).

42. Roncero, C. & Casadaban, M. J. Genetic analysis of the genes involved in synthesis of the lipopolysaccharide core in Escherichia coli K-12: three operons in the rfa locus. Journal of Bacteriology 174, 3250 (1992).

43. Cannistrato, V. J. & Kennell, D. RNase I*, a form of RNase I, and mRNA degradation in Escherichia coli. Journal of Bacteriology 173, 4653 (1991).

44. Hawkins, J. S., Wong, S., Peters, J. M., Almeida, R. & Qi, L. S. Targeted Transcriptional Repression in Bacteria Using CRISPR Interference (CRISPRi). Methods in molecular biology (Clifton, N.J.) 1311, 349–362 (2015).

45. Nomura, M., Gourse, R. & Baughman, G. REGULATION OF THE SYNTHESIS OF RIBOSOMES AND RIBOSOMAL COMPONENTS. Annual Reviews of Biochemistry 53, 75–117 (2003).

46. Andersen, J. B. et al. New unstable variants of green fluorescent protein for studies of transient gene expression in bacteria. Applied and Environmental Microbiology (1998).

47. Akiyama, T. et al. Resuscitation of Pseudomonas aeruginosa from dormancy requires hibernation promoting factor (PA4463) for ribosome preservation. Proceedings of the National Academy of Sciences of the United States of America 114, 3204–3209 (2017).

48. Hauryliuk, V., Atkinson, G. C., Murakami, K. S., Tenson, T. & Gerdes, K. Recent functional insights into the role of (p)ppGpp in bacterial physiology HHS Public Access. Nat Rev Microbiol 13, 298–309 (2015).

49. Porwollik, S. et al. Defined Single-Gene and Multi-Gene Deletion Mutant Collections in Salmonella enterica sv Typhimurium. PLoS ONE 9 (ed Hensel, M.) e99820 (2014).

50. Datsenko, K. A. & Wanner, B. L. One-step inactivation of chromosomal genes in Escherichia coli K-12 using PCR products. Proceedings of the National Academy of Sciences of the United States of America (2000).

51. Labun, K. et al. CHOPCHOP v3: expanding the CRISPR web toolbox beyond genome editing. Nucleic Acids Research 47, W171–W174 (2019).

52. Ferrières, L. et al. Silent mischief: Bacteriophage Mu insertions contaminate products of Escherichia coli random mutagenesis performed using suicidal transposon delivery plasmids mobilized by broad-host-range RP4 conjugative machinery. Journal of Bacteriology 192, 6418–6427 (2010).

53. Peters, J. M. et al. Enabling genetic analysis of diverse bacteria with Mobile-CRISPRi. Nature Microbiology 2019 4:2 4, 244–250 (2019).

54. Neidhardt, F. C., Bloch, P. L. & Smith, D. F. Culture Medium for Enterobacteria. Journal of Bacteriology 119, 736–747 (1974).

55. Berg, S. et al. ilastik: interactive machine learning for (bio)image analysis. Nature Methods (2019).

56. Hatzenpichler, R. et al. In situ visualization of newly synthesized proteins in environmental microbes using amino acid tagging and click chemistry. Environmental Microbiology 16, 2568 (2014).

57. Hall, B. G., Acar, H., Nandipati, A. & Barlow, M. Growth Rates Made Easy. Molecular Biology and Evolution 31, 232–238 (2014).

58. Schwartzman, O. et al. UMI-4C for quantitative and targeted chromosomal contact profiling. Nature Methods 2016 13:8 13, 685–691 (2016).

59. Love, M. I., Huber, W. & Anders, S. Moderated estimation of fold change and dispersion for RNA-seq data with DESeq2. Genome Biology 15, 1–21 (2014).

## References

1. Lin, J. & Amir, A. Homeostasis of protein and mRNA concentrations in growing cells. Nature Communications 2018 9:1 9, 1–11 (2018).

2. Basan, M. et al. Inflating bacterial cells by increased protein synthesis. Molecular Systems Biology 11, 836 (2015).

3. Kubitschek, H. E., Baldwin, W. W., Schroeter, S. J. & Graetzer, R. Independence of buoyant cell density and growth rate in Escherichia coli. Journal of Bacteriology 158, 296–299 (1984).

4. Rollin, R., Joanny, J.-F. & Sens, P. Cell size scaling laws: a unified theory. bioRxiv, 2022.08.01.502021 (2022).

5. Scott, M., Gunderson, C. W., Mateescu, E. M., Zhang, Z. & Hwa, T. Interdependence of Cell Growth and Gene Expression: Origins and Consequences. Science 330, 1099–1102 (2010).

6. Prossliner, T., Gerdes, K., Sørensen, M. A. & Winther, K. S. Hibernation factors directly block ribonucleases from entering the ribosome in response to starvation. Nucleic Acids Research 49, 2226–2239 (2021).

7. Elowitz, M. B., Levine, A. J., Siggia, E. D. & Swain, P. S. Stochastic gene expression in a single cell. Science 297, 1183–1186 (2002).

8. Levien, E., Min, J., Kondev, J. & Amir, A. Non-genetic variability in microbial populations: survival strategy or nuisance? Reports on Progress in Physics 84, 116601 (2021).

9. Eldar, A. & Elowitz, M. B. Functional roles for noise in genetic circuits. Nature 2010 467:7312 467, 167–173 (2010).

10. Lin, J. & Amir, A. Disentangling Intrinsic and Extrinsic Gene Expression Noise in Growing Cells. Physical Review Letters 126, 078101 (2021).

11. Kumar, V. et al. Speckle noise reduction strategies in laser-based projection imaging, fluorescence microscopy, and digital holography with uniform illumination, improved image sharpness, and resolution. Optics & Laser Technology 141, 107079 (2021).

12. Bioucas-Dias, J. M. & Figueiredo, M. A. Multiplicative noise removal using variable splitting and constrained optimization. IEEE Transactions on Image Processing 19, 1720–1730 (2010).

13. Murphy, E. A. One cause? Many causes?: The argument from the bimodal distribution. Journal of Chronic Diseases 17, 301–324 (1964).

14. Veening, J.-W., Smits, W. K. & Kuipers, O. P. Bistability, Epigenetics, and Bet-Hedging in Bacteria. Annual Review of Microbiology 62, 193–210 (2008).

